# Systematic Analysis of Targets of Pumilio (PUM)-Mediated mRNA Decay Identifies a Role of PUM1 in Regulating DNA Damage Response Pathway

**DOI:** 10.1101/387381

**Authors:** Toshimichi Yamada, Naoto Imamachi, Katsutoshi Imamura, Takeshi Kawamura, Yutaka Suzuki, Masami Nagahama, Nobuyoshi Akimitsu

## Abstract

RNA-binding proteins (RBPs) play a pivotal role in gene expression by modulating the stability of transcripts; however, the identification of degradation targets of RBPs remains difficult. Here, we identified 48 target mRNAs of human Pumilio 1 (PUM1), an evolutionally conserved RBP, by combined analysis of transcriptome-wide mRNA stabilities and the binding of mRNAs to PUM1. Here, we developed an approach to identify mRNA targets of Pumilio 1 (PUM1), an evolutionally conserved RBP. By combined analysis of transcriptome-wide mRNA stabilities and the binding of mRNAs to PUM1, we identified 48 mRNAs that both bound to PUM1 and exhibited PUM1-dependent degradation. Analysis of changes in the abundance of PUM1 and its targets in RNA-seq data indicated that DNA-damaging agents negatively regulated PUM1-mediated mRNA decay. Cells exposed to cisplatin had reduced PUM1 abundance and increased *PCNA* and *UBE2A* mRNAs, encoding proteins involved in DNA repair by translesion synthesis (TLS). Cells overexpressing PUM1 exhibited impaired DNA synthesis and TLS and increased sensitivity to the cytotoxic effect of cisplatin. Thus, our method identified targets of PUM1-mediated decay and revealed that cells respond to DNA damage by inhibiting PUM1-mediated mRNA decay to activate TLS.

## Introduction

The ability of cells to sense and adapt to environmental changes is critical for cell survival^1^. Cells modify not only transcription but also mRNA decay rate to change the abundance of mRNAs in respond to external cues^2, 3^. In mammalian cells, mRNA stability is tuned by regulating the activity of decay machinery^4^. Cytoplasmic decay factors are mainly categorized into two categories: ribonucleases and RNA-binding proteins (RBPs). A representative ribonuclease is the CCR4-NOT complex, which contains deadenylating enzymes and triggers exonuclease-mediated decay by shortening poly(A) tail^5, 6^. Consequently, transcripts with the shortened poly(A) tail are degraded by other ribonucleases, such as XRN1 or the RNA exosome complex. Most ribonucleases are targeted to their mRNA substrates through RBPs that specifically bind the target mRNAs and the ribonuclease through physical interactions^7^. Hence, RBPs play a central role in gene regulation by providing specificity to mRNA degradation. Studies that applied high-throughput sequencing methods, such as RIP-seq or CLIP-seq^8–10^, revealed that an individual RBP can bind to 100 to 1,000 mRNAs. For the RBPs involved in mRNA decay, not all mRNAs that bind the RBP are targets of degradation. For instance, only subset of the mRNAs that bind to UPF1, an RBP involved in the RNA degradation and RNA quality control, is subject to UPF1-dependent mRNA degradation^11^. Consequently, knowledge of the mRNAs that bind an RBP is insufficient. This information must be combined with data on mRNA stability to determine the functional targets of an RBP involved in mRNA decay.

Pumilio (PUM) is an evolutionally conserved family of RBPs that regulate the abundance of specific mRNAs and contribute to development, neuronal function, and fertility^12, 13^. For example, *Drosophila melanogaster* Pumilio promotes the degradation of *hunchback* mRNA to control embryonic development and promotes the degradation of *Cyclin B* and *Mei-P26* mRNAs to maintain stem cell self-renewal^14, 15^. Mammals have two PUM genes, PUM1 and PUM2. Mouse and human PUM1, one of two PUM-encoding mammalian genes, promotes the degradation of *ATXN1* mRNA, and haploinsufficiency or complete knockout of *PUM1* in mice leads to neurodegeneration and impaired motor function similar to SCA1 (spinocerebellar ataxia type 1)^16^. A study in rats showed that PUM2 is also important in nerves by maintaining the proper amount of transcripts for the voltage-gated sodium channel Nav1.6^17^. *PUM1* or *PUM2* knockout mice have reduced body mass and impaired fertility^18, 19^. However, PUMs are found ubiquitously in all somatic cells and are not restricted to specific developmental stages or tissues^20, 21^. Therefore, roles for PUMs throughout the body remain to be discovered.

Crystal structure analysis of PUM-RNA complexes showed that RNA-binding domain of PUM (PUM-HD) is composed of eight repeated motifs, and this repeated element specifically recognizes a single 5’-UGUAHAUA-3’ sequence, named PRE (Pumilio Response Element). Several genome-wide measurements of interactions between PUMs and mRNAs verified that PUMs bind to mRNAs with at least one PRE sequence in the 3’UTR^22–24^. PUMs recruit the CCR4-NOT deadenylase complex, resulting in a shortened poly(A) tail, reduced translation, and increased degradation of the target mRNA^25–28^. Because PUMs are excellent model of an RBP to study, extensive prior studies on the interaction between PUMs and mRNAs have been performed ^8, 22, 23, 29, 30^.

Several key questions regarding PUM function remain unanswered. First, what are the complete sets of degradation target mRNAs of PUM1 and PUM2? Although more than 1,000 mRNAs bind to each PUM proteins^22, 23^, those that are subjected to PUM-mediated decay are unknown. Second, what are the ubiquitous biological functions of PUMs in cells? Similar to studies of transcription factors that used large-scale expression analysis in response to chemical perturbations revealed biological functions of the transcription factors^31–34^, we predict that analysis of stimuli that alter PUM-mediated mRNA degradation will reveal their biological functions. However, unlike the existing framework for studying transcription factors, an analytical framework for identification of stimuli that alter RBP-mediated mRNA degradation is limited^35^.

Here, we determined PUM1 function by experimentally identifying PUM1-specific degradation targets and used *in silico* analysis to identify stimuli that modulate PUM-mediated mRNA degradation. We experimentally defined 48 targets of PUM1-mediated decay according to the following criteria: (1) the transcript bound to PUM1 and (2) the stability of the transcript increased in PUM1-depleted cells. In contrast, we found that PUM2 had a little effect on mRNA degradation in the cells examined in this study. *In silico* screening of publicly available RNA-seq data identified DNA-damaging reagents as inhibitors for PUM1-mediated mRNA decay. Subsequently, we found that PUM1 limits the translesion synthesis (TLS) pathway, which enables DNA replication to bypass a site of damaged DNA, and that the DNA-damaging agent cisplatin inhibits this function of PUM1. Thus, we established an approach to identifying biological function of RBPs involved in mRNA decay and discovered a previously uncharacterized role of PUM1 in regulating TLS pathway to repair the damaged DNA present during DNA replication.

## Results

### Identification of target mRNAs of human PUM1 and PUM2 by RIP-seq and BRIC-seq

We defined targets of PUMs by two criteria. One criterion is the mRNA binds to PUMs, which we determined by RNA immunoprecipitation sequencing (RIP-seq)^36^. The other criterion is the mRNA stability is decreased by PUMs, which we determined by 5′-bromo-uridine (BrU) immunoprecipitation chase sequencing (BRIC-seq) ^37, 38^. BRIC-seq enables the determination of genome-wide RNA stability by monitoring the decrease in BrU-labeled RNA abundance over time from which the RNA half-life of each transcript is calculated. By comparing the half-lives of transcripts in control cells with the half-lives of those in PUM-depleted cells, we can identify mRNAs of which stability is regulated by PUMs^11^. Combining the results of the RIP-seq and BRIC-seq analyses yields the targets of PUMs (**Figure 1A**).

**Figure 1.**
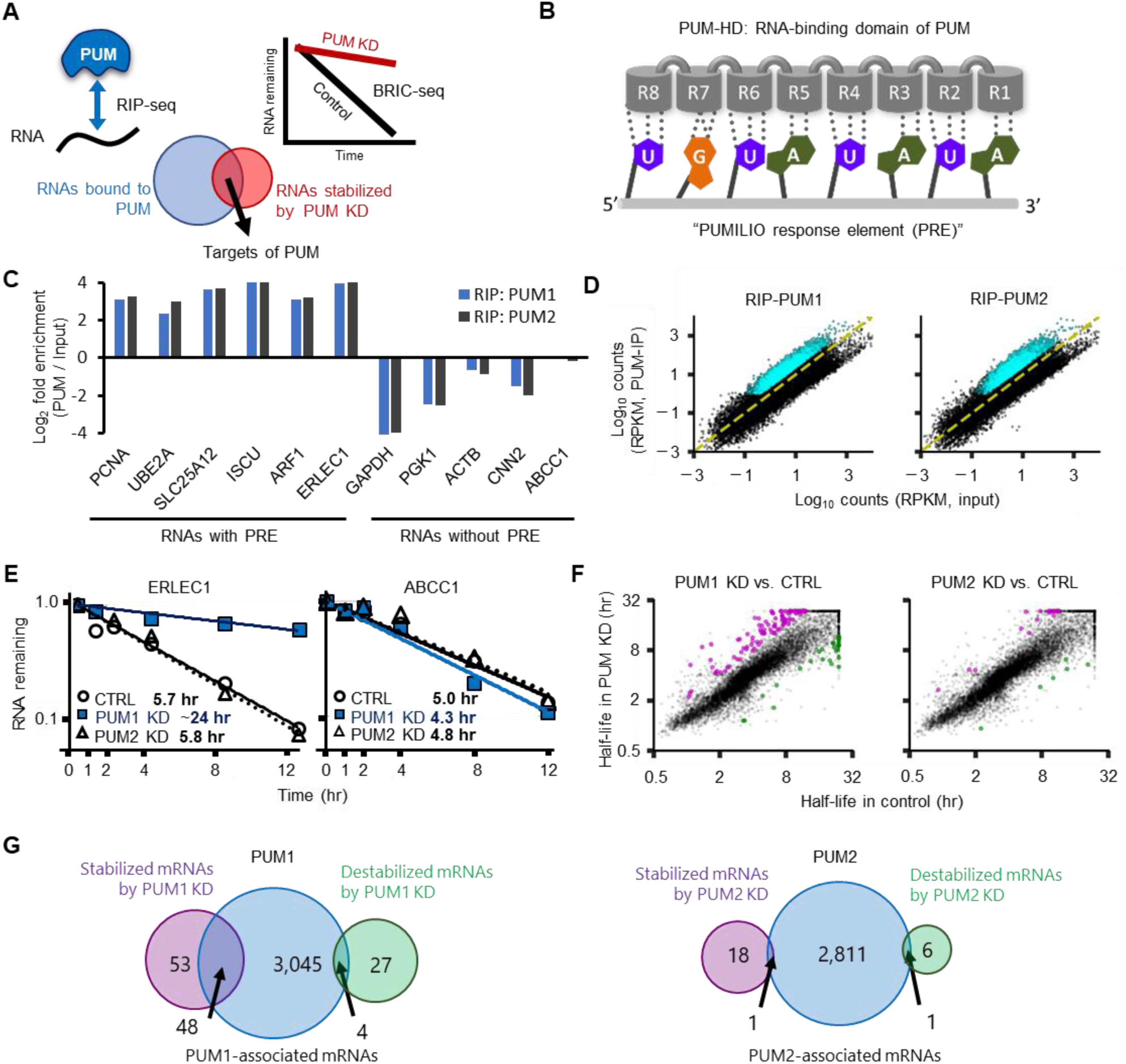
Identification of PUM-target mRNAs. (**A**) A strategy to determine the targets of PUMs. RNAs bound to PUMs were determined by RNA immunoprecipitation sequencing (RIP-seq), whereas RNAs degraded by PUMs were determined by 5′-bromo-uridine (BrU) immunoprecipitation chase-deep sequencing (BRIC-seq). The overlapping RNAs from both analyses are defined as PUM targets in this study. (**B**) mRNA-binding region of human PUM1 and PUM2 proteins. The figure shows the eight repeats (R1 to R8) of the PUM-HD and the specific RNA base of the PUM response element (PRE) that each repeat binds. (**C**) Bar graphs display fold enrichment of the indicated mRNAs representing those with or without PRE. Data were from RIP-seq using FLAG-antibody to immunoprecipitate the proteins from HeLa TO cells. See also figure S1A. (**D**) Scatter plots of RIP-seq data of PUM1 and PUM2. The RPKMs for each transcript identified in the RIP-seq data (RIP-PUM) against the RPKMs for the same transcript in the input sample are plotted. Cyan dots indicate PUM-binding transcripts (FDR < 0.05). See table S1 for transcript identity. (**E**) Representative mRNA decay curves for a PRE-containing transcript (ERLEC1) and a transcript lacking the PRE sequence (ABCC1) under indicated knockdown conditions as determined by BRIC-seq (Figure S1B). In each decay plot, data were fitted with an exponential function. See also Figure S1C, D (**F**) Scatter plots show the mRNA half-life for each transcript calculated by BRIC-seq of PUM1 KD or PUM2 KD cells against the half-life for the same transcript in the control cells. Magenta dots and green dots indicate significantly stabilized transcripts and significantly destabilized transcripts, respectively. See table S1 for list of significantly changing transcripts. (**G**) Venn diagrams show the overlap of the three types of mRNAs: mRNAs for which half-lives were stabilized by knockdown, mRNAs for which half-lives were destabilized by knockdown, and mRNAs that bound to PUM1 or PUM2. The intersection of stabilized mRNAs in PUM1 KD cells and PUM1-associating mRNAs were defined as *bona fide* targets of PUM1-mediated mRNA decay. See also Figure S1 and S2

Human PUM1 and PUM2 share 83% overall similarity with 97% similarity in RNA-binding domain (PUM-HD)^20^, which contains the 8 repeated motifs that interact with a PRE sequence (**Figure 1B**). To identify mRNAs that bind to PUMs, we performed RIP-seq of FLAG-tagged PUM1 or FLAG-tagged PUM2 from HeLa Tet-off (TO) cells expressing similar amounts of either FLAG-tagged protein. As expected, the RIP-seq analysis for each PUM1 and PUM2 showed an enrichment of mRNAs with PRE sequences (**Figure 1C**). With a false discovery rate of 0.05, we found 3,097 and 2,813 mRNAs bound to PUM1 and PUM2, respectively (**Figure 1D, Table S1**). Either PUM1 or PUM2 bound to 3,239 unique transcripts. Of these, 82% bound both PUM proteins. We confirmed the results of RIP-seq using quantitative RT-PCR (RT-qPCR) of selected mRNAs present in the PUM1 RIP or PUM2 RIP (**Figure S1A, upper panel**). In addition, we confirmed a subset of the mRNAs bound to endogenous PUMs by performing RIP with specific antibodies recognizing PUM1 or PUM2 (**Figure S1A, lower panel**).

To investigate the functional classification of the mRNAs associated with PUMs, we searched Protein Analysis THrough Evolutionary Relationship (PANTHER) for biological processes associated with the proteins encoded by the mRNAs that are bound to both PUM1 and PUM2. PANTHER indicated a significant enrichment of components, such as developmental processes (247 mRNAs, FDR < 1.1×10^-4^), mitosis (73 mRNAs, FDR < 1.6×10^-3^), and the MAPK cascade (66 mRNAs, FDR < 1.7×10^-2^) (see **Table S2** for complete results). These three enriched categories are consistent with a previous study of PUM-bound mRNAs ^22^.

To analyze the effect of PUMs on mRNA decay, we performed BRIC-seq to measure changes in mRNA half-lives following the siRNA-mediated knockdown of either PUM1 or PUM2 (**Figure S1B**). By comparing the altered mRNA half-lives between PUM1 and PUM2 knockdown (KD) cells, we found that PUM1 KD had a larger impact on mRNA stability than did PUM2 KD in HeLa TO cells (**Figure 1E, F**). For example, the *ERLEC1* transcripts in PUM1 KD cells were stable for the entire 12 hours of the assay; whereas *ERLEC1* transcripts had a half-life of 5.7 hours in control cells and 5.8 hours in PUM2 KD cells. In contrast, there was no effect of knockdown of either PUM for a transcript that lacked the PRE sequence, such as *ABCC1* mRNA. We validated the BRIC-seq results for 6 transcripts using BRIC followed by RT-qPCR (BRIC-qPCR) (**Figure S1C, S1D**). A genome-wide comparison of the mRNA half-lives indicated that more transcripts were regulated by PUM1 than by PUM2 in these HeLa TO cells (**Figure 1F, Table S1**).

By merging the results of BRIC-seq with those from RIP-seq, we identified 48 transcripts that were both associated with PUM1 and significantly stabilized in PUM1 KD cells (**Figure 1G, Table S1**). In contrast, only four transcripts had shorter half-lives in PUM1 KD cells and were associated with PUM1. Thus, these data indicated that the main role of PUM1 in regulation of mRNA stability in HeLa TO cells is promoting mRNA degradation. For PUM2, only a single transcript belonged in each of the intersections between the RIP-seq and BRIC-seq data (**Figure 1G**); however, neither of these two mRNAs had a PRE sequence. Thus, we did not consider these mRNAs as targets of PUM2.

### Higher abundance of PUM1 explains the dominant effect of PUM1 on mRNA decay

Because PUM1 and PUM2 shared most of their mRNA binding targets (**Figure S2A**) and the enrichment of the PUM1-bound mRNAs was highly correlated with that of the PUM2-bound mRNAs (**Figure S2B**), the binding selectivity of PUM1 and PUM2 for mRNAs appeared highly similar. Therefore, we hypothesized that the negligible effect of PUM2 on mRNA degradation was because PUM1, but not PUM2, preferentially binds to mRNA decay factors. To compare RNA decay factors that interact with PUMs, we isolated PUM1 and PUM2 complexes from cells overexpressing FLAG-tagged PUM1 or FLAG-tagged PUM2 and identified associated proteins by mass spectrometry. Consistent with a previous study in HEK293 cells ^25^, both PUM complexes included multiple subunits of the CCR4-NOT deadenlyase complex, including CNOT1,2,7,9 (**Figure S2C, Table S3**). We confirmed the interaction between FLAG-tagged PUM1 or FLAG-tagged PUM2 and CNOT1 by coimmunoprecipitation followed by Western blotting (**Figure S2D**). We also identified by mass spectrometry other mRNA decay factors, including enhancer of mRNA de-capping (EDC4) and a member of the trinucleotide repeat-containing 6 protein family (TNRC6A)^39–41^. Contrary to our expectation, most of the decay factors had similar yields between PUM1 and PUM2, indicating that both proteins formed similar mRNA decay complexes. The similarity in the mRNA targets and in the mRNA decay factor interactions between both PUMs indicated that PUM2 also has the potential to promote mRNA degradation.

To explain the difference in PUM knockdown on mRNA stability, we estimated the abundance of endogenous PUM1 and PUM2 by quantitative Western blot (**Figure S2E**). By normalizing the PUM band intensity with the GAPDH band and FLAG-tag band intensities, the endogenous PUM1/PUM2 ratio was estimated as 7.1 ± 0.1 (**Figure S2F**). This ratio is similar to the value quantified for HCT116 cells (PUM1/PUM2 = 7.5)^42^. Based on this measurement, we propose that the difference in protein abundance may explain why PUM2 KD does not affect mRNA stability in HeLa TO cells, despite PUM2 exhibiting similar binding ability for mRNAs and RNA decay factors as that of PUM1 (**Figure S2B, S2C**). If both PUM1 and PUM2 have the same potential to control mRNA stability in HeLa TO cells, even complete loss of PUM2 by RNA silencing only results in a 13% reduction in total PUM protein, which would enable the remaining PUM proteins (PUM1) to degrade mRNAs with similar kinetics as those observed in the control condition (**Figure S2G**). Thus, the negligible effect of PUM2 on mRNA degradation in HeLa TO cells may be explained by the low abundance of PUM2.

### *In silico* screening identifies DNA-damaging reagents as inhibitors of PUM1-mediated mRNA decay

The bioinformatic analysis using PANTHER did not show any functional enrichment in the 48 PUM1 targets, which suggests that a novel analytical pipeline is required to investigate the biological role(s) of PUM1-mediated mRNA decay. To this end, we developed an *in silico* screening scheme to identify stimuli that influence PUM1-mediated mRNA decay. For our analysis, we assumed that a relevant stimulus causes inversely correlated changes between PUM1 protein or transcript abundance and the abundance of its target mRNAs: Stimulus that decreases PUM1 abundance should increase the abundance of PUM1 target mRNAs and vice versa (**Figure 2A**). For this *in silico* screening, we collected 491 RNA-seq data obtained in various stress conditions/stimuli from Gene Expression Omnibus (GEO, **Table S2**). To quantify whether the overall trend of PUM1 targets is increased or decreased following a stimulus, we introduced a “variation index,” which we defined by the difference between the number of increased and the number of decreased PUM1 targets relative to the control condition (**Methods**). We plotted the variation index for PUM1 targets in each condition against the corresponding log_2_-fold-change in *PUM1* mRNA to identify those stimuli that promote or inhibit PUM1-mediated mRNA decay (**Figure 2B**).

**Figure 2.**
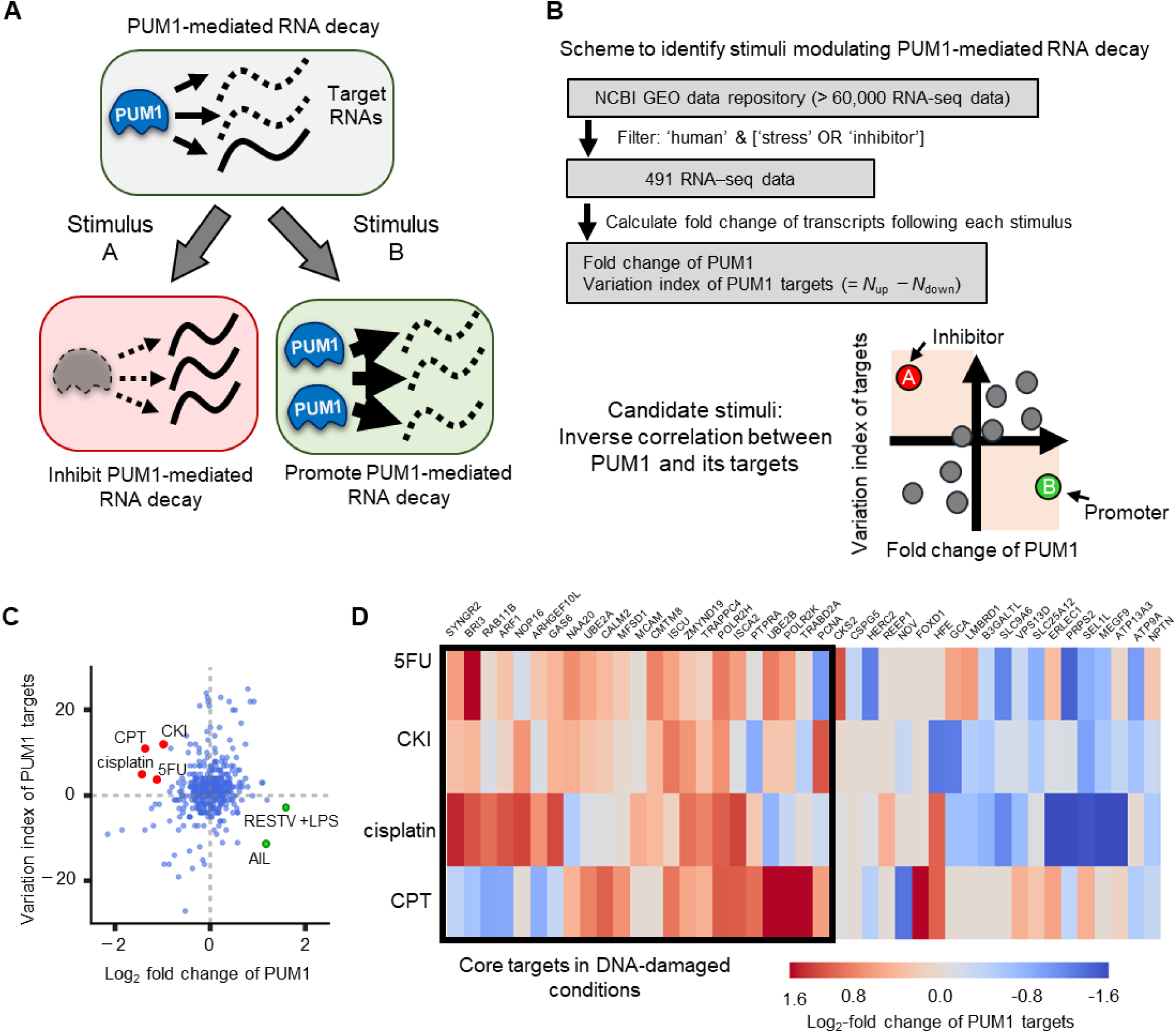
Identification of stimuli that affect PUM1-mediated mRNA decay. (**A**) The concept of the *in silico* screen. The stimuli that inversely change the abundance of PUM1 and PUM1 targets are candidate modulators of PUM1-mediated mRNA decay. (**B**) Analysis pipeline to identify stimuli that modulate PUM1-mediated mRNA decay. The fold change of *PUM1* mRNA and the variation index [(number of upregulated PUM1 targets) – (number of downregulated PUM1 targets)] were calculated for each condition and then plotted to identify potential inhibitors and promoters of PUM1-mediated RNA decay. (**C**) Log_2_-fold change of PUM1 versus variation index for 491 stimuli. Candidate inhibitory stimuli are shown as red dots; candidate promoting stimuli are shown as green dots. See also Figure S3. (**D**) Heat map of fold-change in the PUM1 targets (48 genes) RNA-seq data from cells exposed to the indicated stimuli. Core targets are defined based by hierarchy clustering analysis in which core targets exhibit similar behavior, consistent with stabilization, upon DNA-damaging reagents (see method for details of clustering criteria).

Using this strategy, we identified both stimuli that potentially promoted PUM1-mediated mRNA decay and stimuli that potentially inhibited this process. We found that ailanthone (AIL) and a mixture of Reston virus (RESTV) and lipopolysaccharide (LPS) were potential promoters of PUM1-mediated mRNA decay (**Figure 2C**). In contrast, we found that well-known anti-cancer drugs, such as fluorouracil (5-FU), camptothecin (CPT), and cisplatin, which cause DNA damage and induce apoptosis by inhibiting DNA replication or RNA transcription or both, were potential inhibitors of PUM1-mediated mRNA decay (**Figure 2C**). We also identified Compound Kushen Injection (CKI), a mixture of natural compounds extracted from Kuehen and Baituling in China, as a potential inhibitor of PUM1-mediated mRNA decay. Although the molecular mechanism of CKI remains under investigation ^43^, similarities in the cellular transcriptional response between CKI and 5-FU suggest that CKI also causes DNA damage ^44^. Thus, our *in silico* screening of RNA-seq data obtained under various conditions/stimulus suggested that DNA damage suppresses PUM1-mediated mRNA decay.

Notably, when we calculated the variation index using all 3,097 mRNAs that bound to PUM1 instead of the 48 mRNA-decay targets, *in silico* screening failed to detect these DNA-damaging reagents as potential inhibitors of PUM1-mediated mRNA decay (**Figure S3A**). This failure showed, even though the dataset is larger, analysis that relies only the interaction between PUM1 and mRNAs cannot identify stimuli that modulate PUM1-mediated mRNA decay. Furthermore, we determined that using only the change in *PUM1* transcripts is insufficient to identify DNA-damaging reagents collectively as inhibitors, because the relatively small decrease in *PUM1* in 5-FU– or CKI–treated cells would not identify these DNA-damaging reagents with camptothecin and cisplatin as top-ranking stimuli that altered PUM1 activity (**Figure S3B**).

Not all 48 PUM1 mRNA targets increased in response to DNA-damaging reagents (**Figure 2D**). We expect that this is because DNA-damaging reagents affect both mRNA decay and transcription, with the net response representing the balance between these two effects for any particular transcript. Of the 48 PUM1 mRNA targets, we found 23 transcripts that were increased in at least two of the DNA-damaging conditions and did not exhibit a major decrease (log_2_-fold-change < -1.0) in any condition by hierarchy clustering. Thus, we extracted from the 48 PUM1 targets 23 ‘core targets’, for which the transcript abundance was consistently increased after DNA damage (**Figure 2D**). To further investigate the biological role of PUM1-mediated mRNA decay in cells exposed to DNA-damaging agents, we focused on this set of core targets.

### Suppression of PUM1-mediated mRNA decay by DNA-damaging reagents

We verified the results of the *in silico* screening by measuring the amount of PUM1 and the amounts of selected target mRNAs in drug-treated cells. To select the three core target mRNAs for detailed analysis and verification of the screening results, we chose two genes that are related to DNA repair after DNA damage (*PCNA* and *UBE2A*) and randomly picked up a representative gene (*ISCU*). We also evaluated one non-core target, *SLC25A12* mRNA. By analyzing both core and non-core targets, we could confirm that the core targets represented functionally relevant targets of PUM1-mediated mRNA decay. We analyzed the abundance of these four mRNAs and that of *PUM1* using RT-qPCR in HeLa TO cells under control conditions and in response to exposure to cisplatin or CPT. Corresponding to the RNA-seq data used for the *in silico* screen, cisplatin decreased both the mRNA and protein abundance of PUM1 (**Figure 3A**), whereas the abundance of all four PUM1-target transcripts was increased in HeLa TO cells (**Figure 3B**). As expected, the PUM1-target mRNAs were stabilized in cisplatin-treated cells (**Figure 3C**). Inverse correlations between PUM1 and the core targets also occurred in human colorectal carcinoma (HCT116) and human adenocarcinoma (A549) cell lines (**Figure S4**). Consistent with *SLC25A12* mRNA as a non-core PUM1 target, the abundance of this mRNA did not change in response to cisplatin in HCT116 or A549 cells. In HeLa TO cells, CPT decreased mRNA and protein abundance of PUM1 (**Figure 3A**) and increased only the core PUM1-target mRNA levels (**Figure 3B**).

**Figure 3.**
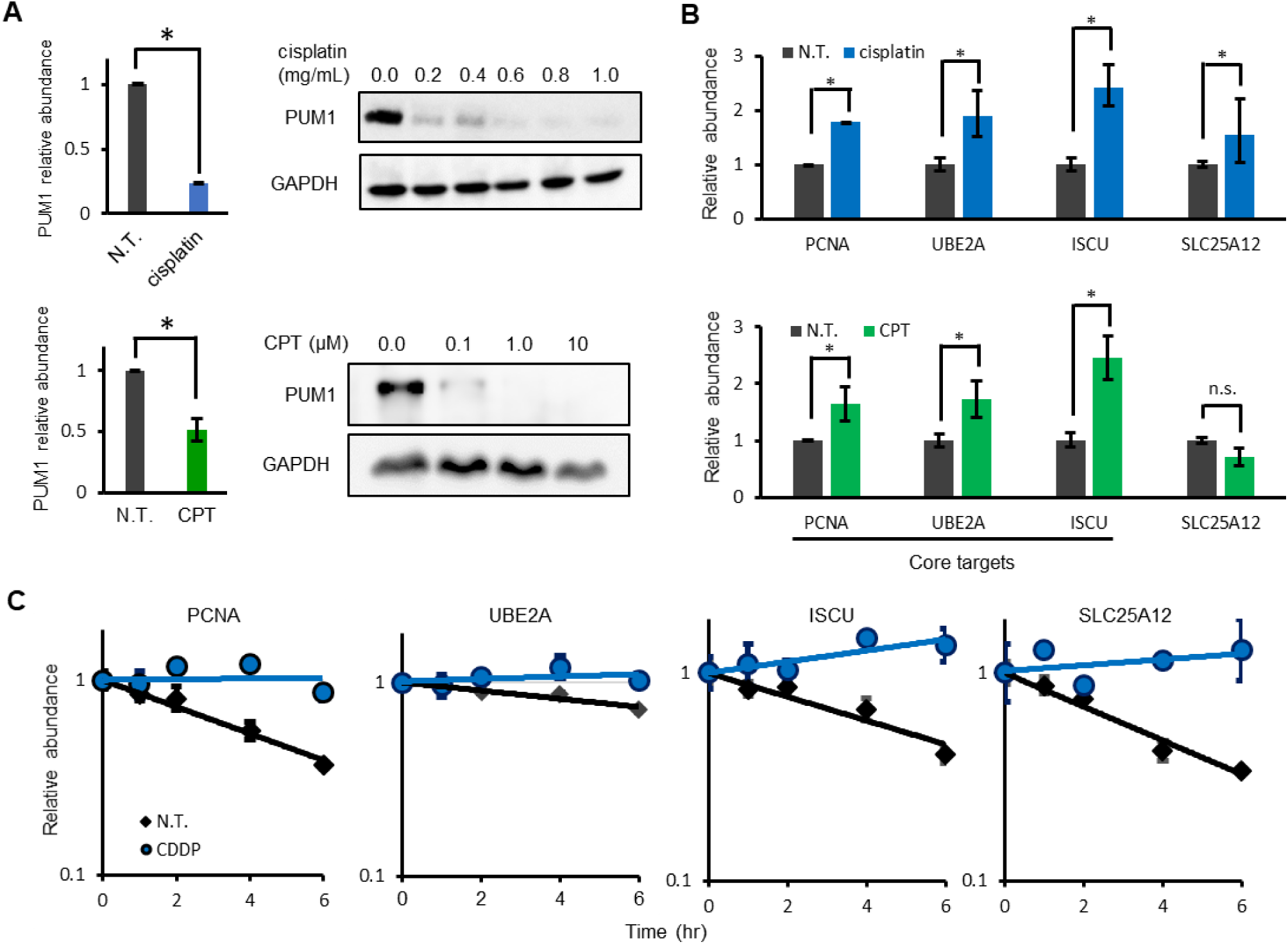
DNA-damaging agents inhibit PUM1-mediated mRNA decay. (**A**) Effect of cisplatin or camptothecin (CPT) on PUM1 mRNA and protein abundance in HeLa TO cells. Cells were exposed to cisplatin (0.5 mg/mL) or CPT (0.1 μM) for 6 hr. PUM1 mRNA was quantified by RT-qPCR; PUM1 protein was analyzed by Western blot. Western blot data are representative of three experiments. (**B**) Changes in the abundance of the indicated PUM1-target mRNAs in non-treated HeLa cells or cells treated with cisplatin (0.5 mg/mL) or CPT (0.1 μM) for 6 hr. (**C**) Time course of abundance of the indicated transcripts in HeLa cells treated with or without cisplatin. Total RNAs were isolated at indicated time points after actinomycin D treatment and relative mRNA abundances (log_10_-transfomed) over time are plotted. For quantification of all transcript abundance data, transcript amounts were determined by Real-time RT-qPCR normalized to geometric mean of *GAPDH* and *PGK1* mRNAs and are shown as mean ± SD. *: p value < 0.05 (Student’s t-test, n=3). The amount in non-treated cells (A and B) or in non-treated cells at time 0 (C) was set to 1. N. T., not treated with either agent; CPT or, cisplatin. See also Figure S4.

Collectively, these results support the *in silico* findings that the DNA-damaging reagents decreased PUM1 transcripts and showed that this decrease results in a reduction in PUM1 protein abundance. These results also support *in silico* finding that PUM1 regulates the PUM1 targets by affecting limiting transcript abundance and showed that the mechanism was by reducing target mRNA stability. Finally, these results support the classification of core and non-core PUM1 mRNA targets: The core target mRNAs— *PCNA*, *UBE2A*, and *ISCU—*were consistently increased by Stimuli that reduced PUM1 abundance; whereas the increase in the non-core target, *SLC25A12*, mRNA depended on the cell type. Thus, the experimental data indicated that the core mRNA targets were tightly and consistently regulated by PUM1-mediated mRNA decay in multiple cell types.

To investigate whether a cause-and-effect relationship exists between the decrease in PUM1 and the increase in PUM1-target mRNA abundance, we prepared a HeLa cell line that overexpressed FLAG-tagged PUM1 or a FLAG-tagged PUM1 mutant lacking the RNA-binding domain (ΔPUM-HD). Using RIP (**Figure 4A**), we confirmed that ΔPUM-HD did not bind PUM1-target transcripts (**Figure 4B**).

**Figure 4.**
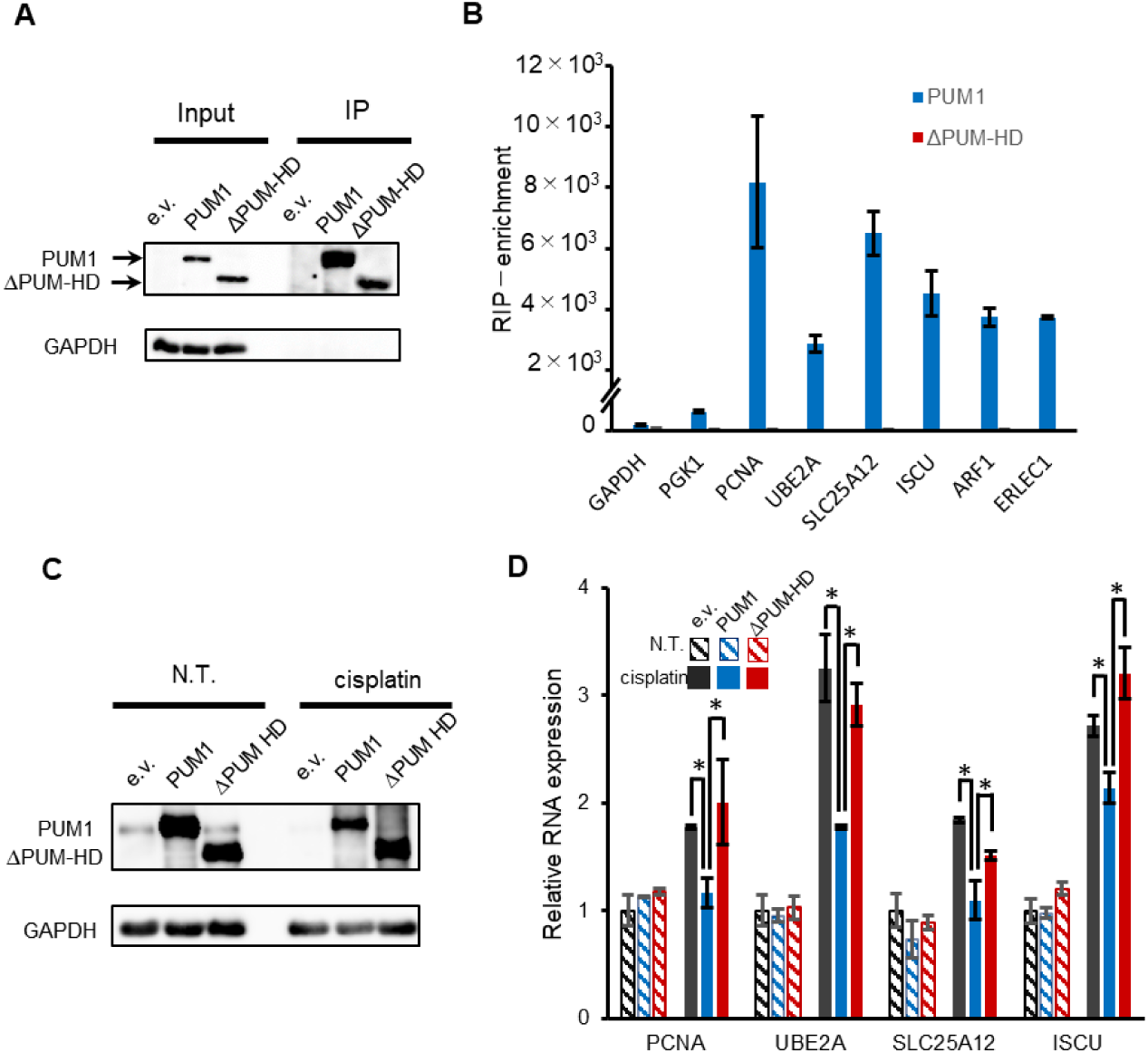
Rescue of DNA damage-induced decrease in endogenous PUM1 by overexpression of FLAG-PUM1. (**A**) Western blot analysis of HeLa cells for exogenous PUM1 in cells transfected with empty vector or in cells overexpressing wild type FLAG-tagged PUM1 or a FLAG-tagged PUM1 mutant lacking the RNA-binding domain (ΔPUM-HD). Samples were used for RIP using a FLAG antibody. Left blot shows a sample used for an RIP experiment. Right blot shows the proteins in the immunoprecipitates from the RIP experiment. The input corresponds to 20% of the total cell lysate (input). GAPDH served as negative control for the immunoprecipitates. (**B**) Analysis of the RNAs bound to PUM1 or ΔPUM-HD. RIP was performed with a FLAG antibody and transcripts quantified by real-time RT-qPCR. Enrichment relative to the amount immunoprecipitated from cells transfected with empty vector are shown. (**C**) Abundance of PUM1 in HeLa cells treated with or without cisplatin. Cells were transfected with empty vector (e.v.), wild-type PUM1, or ΔPUM-HD. For the treated condition, cells were exposed to cisplatin (0.5 μg/mL) for 6 hr. Samples from non-treated (N.T.) and treated (cisplatin) cells were used for protein analysis by Western blot using an antibody recognizing PUM1 or mRNA analysis. GAPDH served as a loading control. (**D**) The effect of PUM1 overexpression on the abundance of PUM1 targets in cells with DNA damage. Real-time RT-qPCR was performed for the indicated PUM1 targets from samples like those shown in panel C. The abundance of PUM1-target transcripts was normalized to the geometric mean of *GAPDH* and *PGK1* mRNAs in e.v.-transfected cells with N.T. condition set to 1. Data are shown as mean ± SD (n=3). *: *p* value < 0.05 (Student’s *t*-test).

In the overexpressing cell lines, FLAG-PUM1 or FLAG-ΔPUM-HD was abundant in cells exposed to cisplatin (**Figure 4C**). We analyzed the abundances of mRNA for four PUM1 targets in the control cells with the empty vector and in the FLAG-PUM1– or FLAG-ΔPUM-HD–overexpressing cells (**Figure 4D**). Compared with any of the non-treated cells, cells overexpressing FLAG-PUM1 did not exhibit increases in the amounts of *PCNA* and *SLC25A12* mRNAs in response to cisplatin treatment, indicating that these transcripts were primarily regulated by PUM1. Compared with the amounts in the non-treated cells, *UBE2A* and *ISCU* mRNAs were increased by cisplatin, indicating that FLAG-PUM1 is partially responsible for the regulation of these transcripts. Compared with the amounts in the cisplatin-treated cells overexpressing FLAG-PUM1, the amounts of the mRNAs of all four targets were significantly increased in the cells overexpressing FLAG-ΔPUM-HD or the control cells with the empty vector after cisplatin treatment. These results showed that DNA damaged-induced increase in some PUM1 targets (represented by *PCNA* and *SLC25A12*) strictly depended on a reduction in functional PUM1. Furthermore, RNA-binding was required for PUM1-mediated effects on mRNA abundance in response to DNA damage, consistent with changes in PUM1-mediated mRNA decay contributing to the increases in mRNA abundance.

### Activation of translesion synthesis after DNA damage by suppression of PUM1-mediated mRNA decay

Platinum-based drugs, such as cisplatin, induce DNA interstrand crosslinks (ICLs)^45, 46^. If cells fail to repair the damaged DNA prior to DNA replication in S phase, DNA polymerase stalls at the site of damaged DNA during replication. A critical process in the repair of an ICL present during DNA replication is TLS, which involves DNA synthesis by TLS polymerases, such as Pol η, to bypass the damaged template^47–49^. UBE2A (also known as RAD6A) and PCNA are fundamental components in TLS^50, 51^. Therefore, we hypothesized that cells activate TLS by suppressing PUM1-mediated mRNA decay.

PCNA mono-ubiquitination by UBE2A triggers TLS^52, 53^. To examine whether suppression of PUM1 increased mono-ubiquitination of PCNA, HCT116 cells were exposed to cisplatin. Cisplatin increased the abundance of UBE2A and PCNA, as well as the amount of mono-ubiquitinated PCNA (**Figure 5A**). Consistent with these increases involving a decrease in PUM1-mediated mRNA decay, PUM1 overexpression attenuated the cisplatin-induced increases in the UBE2A and PCNA and the mono-ubiquitination of PCNA.

**Figure 5.**
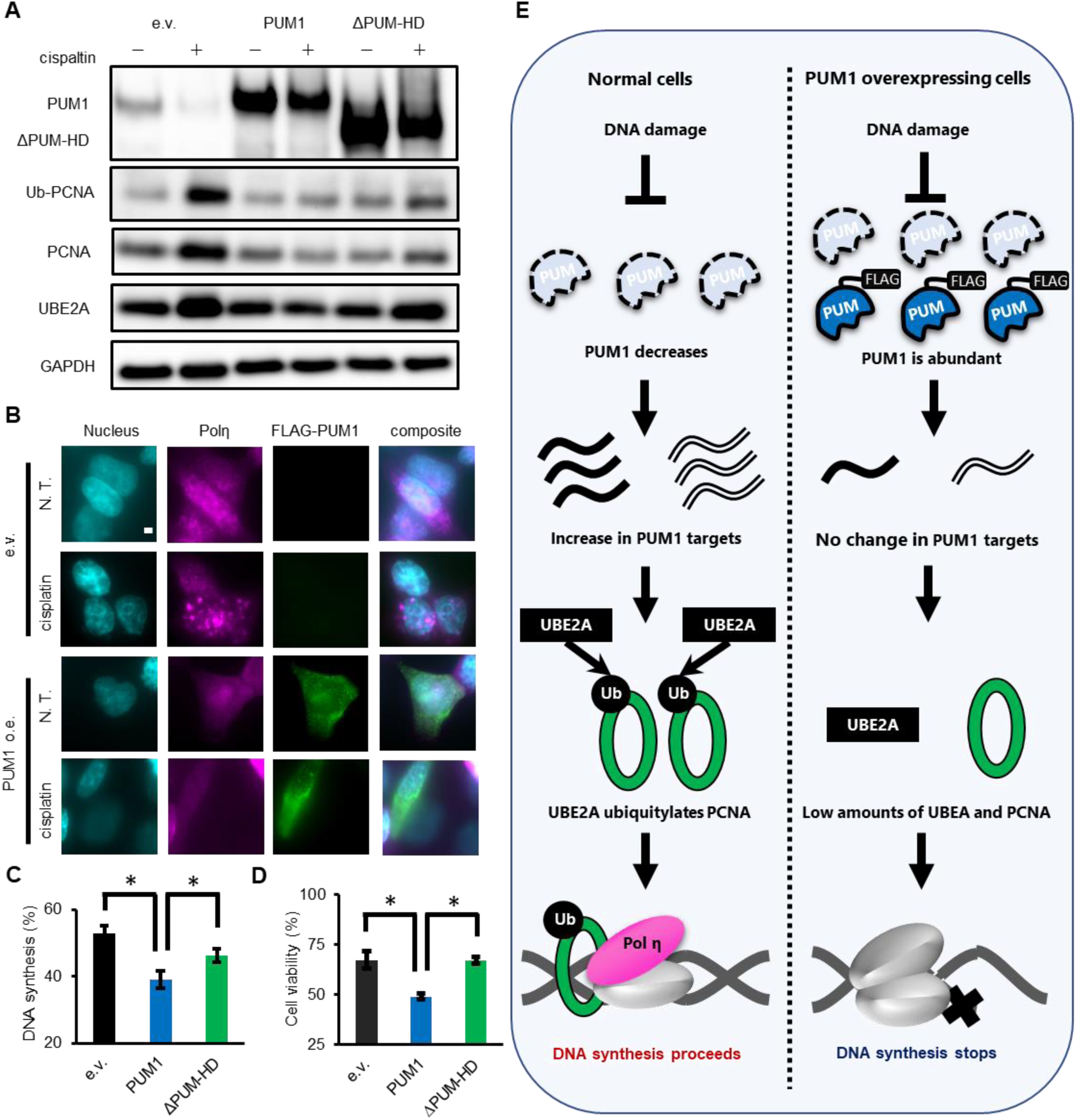
Cells activate translesion synthesis by suppressing PUM1-mediated mRNA decay. (**A**) Effect of cisplatin on the abundance of PUM1 targets encoding proteins involved in TLS. The indicated proteins in control HCT116 cells transfected with empty vector (e.v.), FLAG-tagged wild-type PUM1, or FLAG-tagged ΔPUM-HD with or without cisplatin exposure (0.5 mg/mL for 6 hr). Data are representative of two experiments. (**B**) Effect of cisplatin on the distribution of Pol η and FLAG-tagged PUM1 in HCT116 cells with or without cisplatin exposure (0.5 mg/mL for 6 hr). Immunofluorescence staining of Pol η (magenta) and FLAG-PUM1 (detected with FLAG antibody, green) in cells counterstained with DAPI (cyan). Images are representatives of 40-80 cells (see also Figure S5B) (**C**, **D**) Percent of cells with active DNA synthesis (C) or percent of viable cells (D) in cisplatin-treated (0.5 mg/mL for 6 hr) HCT116 cells transiently transfected with empty vector (e.v.), FLAG-PUM1, or FLAG-ΔPUM-HD. *: *p* value < 0.05 (Student’s *t*-test, n=3). (**E**) Model of PUM1 function in activation of TLS after DNA damage.

As additional evidence of activation of TLS involving a reduction in PUM1-mediated mRNA decay, we analyzed the distribution of Pol η in response to cisplatin in control HCT116 cells and in cells overexpressing FLAG-tagged PUM1. Pol η accumulates at stalled replication forks^54^. We expected that cells with compromised TLS would exhibit a difference in Pol η distribution after the induction of DNA damage. Pol η abundance was similar in cells that had or had not been exposed to cisplatin (**Figure S5A**). In the absence of exposure to a DNA-damaging agent, Pol η was homogenously distributed in nucleus. Cisplatin treatment induced Pol η foci formation; however, fewer cells formed Pol η foci when PUM1 was overexpressed (**Figure 5B, S5B**), indicating that the decrease in PUM1 plays an important role in the localization of Pol η in response to unrepaired DNA damage.

The presence of Pol η foci is believed to indicate sites of DNA synthesis^55, 56^. Thus, we predicated that, by inhibiting TLS, PUM1 overexpression would impair DNA synthesis. To quantify DNA synthesis, we performed BrdU incorporation assays. Relative to cells that had not been exposed to a DNA-damaging agent, cisplatin reduced DNA synthesis (**Figure 5C**). Overexpressing PUM1, but not ΔPUM-HD, significantly reduced DNA synthesis to 39% in response to cisplatin, which was significantly lower than the amount in normal cells exposed to cisplatin (53%). These data indicated that PUM1 overexpression inhibits TLS and impairs DNA synthesis in cells exposed to a DNA-damaging agent.

Inhibition of TLS in cells with high amounts of DNA damage induces cell death^57–59^. Thus, we hypothesized that cells acquire resistance to death induced by cisplatin by suppressing PUM1-mediated mRNA decay. To test this hypothesis, we quantified the cisplatin-induced differences in viability of cells that overexpressed PUM1 or ΔPUM-HD. Compared with control cells exposed to cisplatin, PUM1 overexpressing cell viability was significantly lower (**Figure 5D**). In contrast, the overexpression of ΔPUM-HD, which did not restore PUM-mediated mRNA decay, did not reduce the viability of cells exposed to cisplatin (**Figure 5D**). Taken together, our results suggested that the decrease in PUM1 in response to DNA-damaging agents activates the TLS DNA repair pathway and provides a resistance mechanism to the toxicity of such agents (**Figure 5E**).

## Discussion

Here, we identified a previously unknown biological function of PUM1-mediated mRNA decay in regulating the TLS pathway during DDR. We achieved this by developing an approach to determine *bona fide* degradation target mRNAs of PUM1, not just mRNAs that bound PUM1. We used BRIC-seq to determine the mRNAs that are degraded by PUM1 and combined these data with RIP-seq to identify mRNAs that bound PUM1, resulting in the determination of 48 mRNAs that both bind to and are degraded by PUM1 in HeLa cells. We considered these 48 mRNAs *bona fide* degradation target mRNAs of PUM1. Next, by considering PUM1 and its targets a “module”, we sought stimuli that modulate this module, by developing a bioinformatic pipeline for screening the stimuli that altered both *PUM1* transcript abundance and PUM1-mediated mRNA decay (the abundance of the transcripts in the 48 PUM1 target mRNAs). This *in silico* screening revealed that *PUM1* transcripts decreased in cells exposed to one of several DNA-damaging agents and that a subset of transcripts in the PUM1 48-targets was increased. Inspection of the transcripts that increased revealed two mRNAs, *UBE2A* and *PCNA*, that encode proteins mediating DNA repair in response to DNA damage. In experiments with cultured cells, we confirmed that DNA-damaging agents decreased the abundance of PUM1, increased the abundance of UBE2A and PCNA, and activated TLS, which is a critical process for repairing damaged DNA encountered during DNA replication.

Our data fill a key gap in understanding how TLS is activated in response to chemotherapeutic DNA-damaging agents. The DNA ICLs induced by platinum-based drugs^50, 51, 60^, such as cisplatin, activate TLS. UBE2A and UBE2B are ubiquitin ligases that regulate DNA polymerase switching during TLS through mono-ubiquitination of PCNA^52^. Moreover, knocking down or inhibiting either UBE2A or UBE2B attenuated PCNA ubiquitination and decreased activation of DNA repair after cisplatin treatment^51^. Although these studies showed that UBE2A triggers TLS, the activation mechanism of UBE2A was unknown. In this study, we revealed that *UBE2A* mRNA and *PCNA* mRNA are destabilized in cells that do not have extensive ICLs. In response to cisplatin, PUM1 abundance decreased and UBE2A and PCNA, along with monoubiquitinated PCNA, increased. Overexpression of PUM1 inhibited TLS, as indicated by reduced formation of Pol η foci, impaired DNA synthesis, and decreased cell viability. Thus, we found that PUM1-mediated RNA decay is a negative regulator of the TLS and that inhibition of this process contributes to resistance to the toxicity of DNA-damaging agents that cause ICLs, suggesting that activation of this process could increase the effectiveness of such chemotherapeutics.

Little is known about the regulation of mRNA stability during DDR. Short-wavelength ultraviolet light (UVC) induced the stabilization of mRNAs with short half-lives, such as *CDKN1A* (encoding a protein also known as p21) and *FOS* transcripts^61, 62^. Interestingly, CDKN1A protein is also known to negatively regulates TLS by binding to PCNA and, thus, affecting the interaction between PCNA and Pol η ^63, 64^. Although further analysis would be needed, such as genome-wide measurement of mRNA stability after DNA damage, these results suggest that cells adjust mRNA stability to response to DNA damage. From a cell physiology standpoint, reducing mRNA decay and stabilizing transcripts as part of DDR might enable the cell to maintain a pool of normal transcripts that would produce unmutated proteins under damaged DNA. Many DNA lesions arrest transcription, and transcripts expressed from damaged DNA tend to produce abnormal proteins, which could have toxic consequences (**Figure S4C**). Stabilization of the existing mRNAs would be an expedient mechanism to ensure sufficient amounts of undamaged transcripts for cell survival during DDR. Alternatively, the transcripts that are stabilized could be highly specific and tightly regulated, for example by selective control of the abundance or distribution of RBPs, such as the PUMs. Genome-wide measurements of mRNA stability in response to DNA damage will address these questions and provide information to reveal yet-to-be-defined mechanisms that regulate DDR and DNA repair.

The long non-coding RNA (lncRNA) NORAD is proposed to be a regulator of PUM function and genome stability and is increased by DNA damage. NORAD is considered a decoy of PUM, because the sequence of NORAD is comprised of repetitive PUM-binding motifs^65, 66^. NORAD depletion results in aberrant mitosis, and DNA damage induces an increase in the abundance of this lncRNA^42^. Loss of NORAD function impaired proper chromosome segregation even in cells not exposed to DNA-damaging conditions, which was considered related to inhibition of PUM activity^42^. If the abundance of NORAD increases, then PUM-mediated mRNA decay should be inhibited, because NORAD would compete for PUM-target mRNAs. However, another study found NORAD in both the cytoplasm and nucleus, determined that this lncRNA interacts with RBMX in nucleus, a protein involved in DDR, and showed that this NORAD-RBMX interaction maintained genome stability^67^. Furthermore, the NORAD accumulated in the nucleus after DNA damage, suggesting that NORAD binding to PUM, which is a cytoplasmic protein, was not the mechanism of PUM inhibition in cells exposed to DNA-damaging agents. Our study provides a NORAD-independent mechanism for inhibition of PUM proteins in cells with DNA damage: PUM-mediated mRNA decay is inhibited after DNA damage because of decrease in PUM abundance (both transcript and protein), not because of an interaction with NORAD.

*In silico* screening also found that a mixture of Reston virus (RESTV) and lipopolysaccharide (LPS) were potential promoters of PUM1-mediated mRNA decay (**Figure 2C**). The identification of the mixture of RESTV and LPS as promoting PUM1-mediated mRNA decay suggested that infections influence this PUM1-dependent process. This finding is consistent with a previous study showing that PUM proteins positively regulate RIG-I-like receptor signaling, which detect and respond to viral nucleic acids 43. Although the previous study used Newcastle disease virus (NDV), these results suggest that PUM1 has a role in the immune response to infections of diverse pathogens.

Our current analysis indicated a function of PUM1 distinct from those reported by other studies. Functions of PUM1 and its homologous proteins, such as Puf proteins, have been analyzed in various organisms ^12, 24, 68^. In *Saccharomyces cerevisiae*, Puf proteins inhibit cell differentiation and regulate organelle biogenesis ^69^. In *Caenorhabditis elegans*, *Drosophila melanogaster*, and *Xenopus laevis*, PUM proteins regulate several developmental processes, including embryogenesis, gametogenesis and gamete maturation ^70^, and neuronal development ^71^. In mammalian cells, PUM proteins are implicated in the transition from the quiescent to the growth phase ^72^ and in spermatogenesis ^73^. Our *in silico* screen did not detect these functions, but this screen was also not designed to detect such developmentally specific and cell-type specific functions. Consistent with these reported functions, we found many mRNAs that interacted with PUM1 were related to development and cell differentiation by PANTHER pathway analysis (**Table S2**). However, most mRNAs that bind to PUM1 were not destabilized by PUM1-mediated degradation in HeLa TO cells under normal conditions, suggesting that PUM1-mediated degradation may be regulated. For example, posttranslationally modified PUM1 may degrade a certain set of mRNAs ^74, 75^. In growth-factor stimulated cells, PUM1 is phosphorylated and this form mediates *p27* mRNA degradation^72^. This transcript is not listed in our targets of PUM1, because we did not perform experiments to identify targets of PUM1 that depended on PUM1 phosphorylation. To gain a more complete understanding of PUM1 function, it will be necessary to include studies of the posttranslationally modified forms of PUMs. Moreover, to gain detailed insight into the roles of PUM1 in differentiation and mitosis, studies will need to be performed with cells in different stages of differentiation or stem cells or with cells in difference phases of the cell cycle. Each of these can be performed with our analytical pipeline or with adaptation of our pipeline.

Here, we showed that *in silico* screening of RNA-seq data deposited in GEO could predict a previously unknown function of PUM1. Our *in silico* screening is based on the following hypothesis: Conditions that inversely modulate the abundance of *PUM1* transcripts and the abundance of PUM1 targets. We emphasize that the *in silico* screening failed to determine DNA-damaging reagents as inhibitors of PUM1-mediated mRNA decay when the complete set of mRNAs that bound to PUM1 were used for screening (**Figure S3A**). This failure indicates that binding of PUM1 to a transcript is only a necessary condition, but not a sufficient condition, for PUM1 to exhibit a function on the transcripts. Although many studies mainly focus on identifying transcripts bound to the RBPs of interest using RIP-seq or CLIP-seq ^8, 9, 76–78^, our study highlights the importance of identifying *bona-fide* targets of the particular RBP in the analysis of their biological functions.

The capability of *in silico* screening depends on the number and variety of RNA-seq data available in public databases. Currently, our database includes 187 conditions (**Table S4**). Considering that more than 5,000 inhibitors for cell biology are commercially available, the thousands of clinically approved drugs, and the many physiological stimuli, these conditions are only a small fraction of biologically relevant stimuli. We intend to expand our *in silico* screening capability with the addition of RNA-seq data and inclusion of microarray data. Simply including the microarray data in GEO increases the number of data sets meeting search criteria for the keywords ‘inhibitor’ and ‘human’ with ∼400 for RNA-seq and ∼ 1,400 for microarray data. Finally, our approach can be applied to any RBP that regulates mRNA stability. Thus, our method provides a genome-wide view of RBP-mediated mRNA decay and its regulation, which leads to testable hypotheses of RBP functions.

## Acknowledgments

We would like to thank our lab members for the fruitful discussions and help with the experiments. We would like to express our sincere thanks to Prof. Atsushi Shibata for the constructive discussions. We thank Nancy R. Gough, BioSerendipity, LLC for editorial assistance. This work was supported by grants from MEXT KAKENHI (15H04642, 16H06279). T.Y. received the JSPS Research Fellowship for Young Scientists and is supported by a Grant-in-Aid for JSPS Fellows (17J11266).

## Author contributions

Conceptualization, T.Y. and N.A.; Methodology, T.Y., N.I., and K.I.; Software, T.Y., N.I., and S.Y.; Formal Analysis, T.Y. and N.I.; Investigation, T.Y. and T.K.; Resources, T.K. and M.N.; Data Curation, T.Y. and N.I.; Writing – Original Draft, T.Y.; Review & Editing N.I., K.I., T.K., M.N., S.Y., and N.A.; Supervision, N.A.

## Methods

### Contact for Reagent and Resource Sharing

Further information and requests for resources and reagents should be directed to and will be fulfilled by the Lead Contact, Nobuyoshi Akimitsu.

### Experimental Model and Subject Details

#### Cell Culture and Cell Lines

HeLa Tet-off cells (HeLa TO cells) (Clontech, Palp Alto, CA) and A549 cells (kindly provided by Dr. Nobukuni) were grown in Dulbecco’s modified Eagle’s medium (DMEM) (Wako, Osaka, Japan) supplemented with 10% heat-inactivated fetal bovine serum (FBS) (Life Technologies, Grand Island, NY) at 37 °C in a humidified incubator (Thermo Fisher Scientific, Walthman, MA) with 5% CO_2_. HCT116 (obtained from ATCC) cells were maintained in McCoy’s 5A medium (Life Technologies) with 10% FBS (Life Technologies) at 37 °C with 5% CO_2._ All cell lines were banked directly after being purchased from vendors and were used at low passage numbers (less than twenty).

### Method Details

#### Plasmid construction

FLAG-PUM1, FLAG-PUM2, and FLAG-ΔPUM-HD were built using conventional restriction enzyme cloning.

PUM1 and PUM2 cDNA were amplified from a cDNA library. In brief, a first strand cDNA library was synthesized using total RNA isolated from HeLa Tet-off cells and SuperScript II Reverse Transcriptase (Thermo Fisher Scientific). PCR amplification was performed using the following primer sets:

#### For PUM1

Forward: 5’-ccgGATATCagcgttgcatgtgtcttgaag-3’ (Capital letters represent an EcoRV site)

Reverse: 5’-cccccctaatggtatcatctgaGCGGCCGCataagaat-3’ (Capital letters represent a NotI site)

#### For PUM2

Forward: 5’-tgaGAATTCtaatcatgattttcaagctcttgc-3’ (Capital letters represent an EcoRI site)

Reverse: 5’-tgaCTCGAGttacagcattccatttggt-3’ (Capital letters represent an XhoI site)

The PCR product was digested with the corresponding enzymes and inserted into the digested pcDNA 3.1 Hygro vector with a FLAG tag sequence (5’-ACTAGTCCAGTGTGGTGGAATTCTGCA-3’) between BamHI and EcoRV.

To construct the PUM1 mutant, which lacks the RNA-binding domain (ΔPUM-HD), the PUM1 mutant region was amplified from FLAG-PUM1 with the following primer sets:

Forward: 5’-aaaGATATCagcgttgcatgtgtcttg-3’ (Capital letters represent an EcoRV site)

Reverse: 5’-attcttatgGCGGCCGCtcactgctcctgccagaagg-3’ (Capital letters represent a NotI site)

The PCR product was digested with the corresponding enzymes and inserted into the digested pcDNA 3.1 Hygro vector with the FLAG tag sequence.

All plasmids were prepared using the PureYield Plasmid Midiprep System (Promega, Madison, WI) for transfection to cells.

#### Transfection

The plasmids were transfected into HeLa TO cells using Lipofectamine 2000 (Thermo Fisher Scientific) according to the manufacturer’s protocol. In brief, the day prior to the transfection, the cells were plated into 12 wells (for RIP-seq and BRIC-seq, 10 cm dish) with 1.5×10^5^ cells/mL density. For each well, 2 μL of Lipofectamine 2000 reagent mixed with 500 μL of OptiMEM medium (Life Technologies) were added to the 1.0 μg plasmid. After 20 minutes of incubation at room temperature, the mixture was added to the cells, followed by a 6-hour incubation at 37 °C with 5% CO_2_. The OptiMEM medium was subsequently replaced by culturing medium (DMEM with FBS). Transfected cells were used for further studies at 24-hours post-transfection.

All siRNAs were synthesized by Hokkaido System Science (Hokkaido, Japan). siRNAs were transfected into cells using Lipofectamine RNAi MAX (Thermo Fisher Scientific), according to the manufacturer’s protocol. Briefly, siRNA duplexes (final concentration, 10 nM) were transfected into cells, followed by incubation for 6-hours in 37 °C / 5% CO_2_, and the medium was subsequently changed. The cells were then incubated for 48-hours. The siRNA sequences are listed in **Table S5**.

#### Sample preparation for RNA-immunoprecipitation sequencing (RIP-seq)

Plasmid cloned FLAG-tagged PUM1 or PUM2 was transfected into HeLa TO cells (1.5×10^6^ in 10 cm dish) using Lipofectamine 2000 (Thermo Fisher Scientific), as described in the previous section. After 24-hours, the cells were harvested and washed in cold PBS. The cell pellets were lysis by the addition of 300 μL of polysome lysis buffer (PLB: 100 mM KCl, 5 mM MgCl_2_, 10 mM Hepes, pH 7.0, 0.5% NP-40, Recombinant RNase inhibitor (Takara) and protease inhibitor cocktail (Sigma-Aldrich)). The lysate was subsequently sonicated for 2 min with 15 second intervals using a Bioruptor UCD-250 (Cosmo Bio, Tokyo, Japan). After 15 min of centrifugation at 12,000 rpm, 10 μL of supernatants were used for the input protein sample, 30 μL of supernatants were used for the input RNA sample, and the remaining were used for immunoprecipitation (IP).

Antibody-bead conjugates were prepared as follows: One hundred μL of protein G Dyna beads (Thermo Fisher Scientific) were washed three times in PLB and were resuspended in 1 mL of PLB. Fifteen μL of anti-FLAG M2 mouse monoclonal antibody (SIGMA) were added to the beads. After rotating the mixture at 4 °C overnight, the beads were washed with 1 mL PLB two times.

The supernatants were mixed with antibody-bead conjugates and rotated for 2-hours at 4 °C. The beads were washed six times in ice-cold NT2 buffer (50 mM Tris, pH 7.4, 150 mM NaCl, 1 mM MgCl_2_, 0.05% NP-40). After the final wash, the beads were resuspended in 1 mL of NT2 buffer. In 1 mL lysate, 40 μL of lysate were used for the protein sample and the remaining was used for the RNA sample.

For RNA isolation, 200 μL of 100 mM Tris/50 mM NaCl and 600 μL of ISOGEN LS (Nippon Gene, Tokyo, Japan) were subsequently added, followed by RNA isolation according to the manufacturer’s protocol. Prior to sequencing, the isolated RNA was quantified by RT-qPCR, and the size distribution was assessed using an Agilent 2100 Bioanalyzer. The isolated RNA was used for high-throughput sequencing. A RIP-seq complementary DNA library was prepared from 1 µg of total RNA using the TruSeq RNA Library Prep Kit v2 (Illumina, San Diego, CA), according to the manufacturer’s protocol. Thirty-six base pair single-end read RNA-seq tags were generated using an Illumina HiSeq 2500, according to the standard protocol. The fluorescent images were processed to nucleotide sequences using the analysis Pipeline supplied by Illumina.

#### Motif identification

To validate the putative binding motif for PUM in the 3’UTRs of the transcripts bound to PUM, we performed *de novo* motif discovery on the sequences using multiple EM for motif elicitation (MEME). To reduce the calculation time, the top 100 transcripts of the RIP-seq fold change were selected for the MEME analysis. Here, 3’UTR was defined in the RefSeq annotation data.

#### Bromo-uridine (BrU) immunoprecipitation chase-deep sequencing (BRIC-seq)

BRIC was performed as previously described ^38^. In brief, cells were incubated at 37 °C in the presence of 150 μM bromo-uridine (BrU) (Wako Chemical, Tokyo, Japan) for 24-hours in a humidified incubator with 5% CO_2_. Cells were harvested at the indicated time points after replacing BrU-containing medium with BrU-free medium. Total RNA was isolated using RNAiso Plus (Takara), followed by the isolation of BrU-labeled RNA using anti-BrdU mouse antibody (clone 2B1, MBL, Nagoya, Japan). The isolated RNA was used for high-throughput sequencing as described in the “Sample preparation for RNA-immunoprecipitation sequencing (RIP-seq)” section.

#### Sample preparation of IP-Mass spectrometry analysis

The preparation of cell lysate for IP-mass is exactly the same procedure for RIP-seq (refer to the first paragraph of the “Sample preparation for RNA-immunoprecipitation sequencing (RIP-seq)” section). To elute proteins with little antibody contamination, we cross-linked the antibody to the beads. One hundred μL of Dynal protein G beads (Thermo Fisher Scientific) were washed three times in PLB and were resuspended in 1 mL of PLB. Fifteen μL of anti-FLAG M2 mouse monoclonal antibody were added to the beads. After rotating the mixture at 4 °C overnight, the beads were washed two times with 1 mL of 100 mM sodium borate, pH 9.0. The beads were re-suspended with 1 mL DMP/borate solution and were incubated for 15 minutes at room temperature. After the removal of the buffer, the beads were re-suspended with 1 mL fresh DMP/borate solution and were incubated for 30 minutes at room temperature. The beads were subsequently washed two times with 1 mL of 50 mM Glycine, pH 2.5. Finally, the beads were washed two times with PLB and were stored with PLB.

Prior to mixing the beads and cell lysate, cells were treated with 3 µL of RNase A + T1 mixture (1 µg/µL RNaseA and 40 U/µL RNaseT1) for 10 minutes on ice. The cell lysates were mixed with antibody-bead conjugates and rotated for 2-hours at 4 °C. The beads were washed six times in ice-cold NT2 buffer. After the final wash, the beads were resuspended in 20 µL of 0.02% RapiGest to elute the proteins. The beads in RapiGest solution were vortex and were incubated for 30 minutes at 67 °C. After spin down of the beads, the supernatants were used for mass spectrometry.

#### Liquid Chromatography-Tandem Mass Spectrometry (LC/MS/MS)

A capillary reverse phase HPLC-MS/MS system (ZAPLOUS System; AMR Inc.), composed of a Paradigm MS4 quadra solvent delivery device (Michrom BioResources), an HTC PAL autosampler (CTC Analytics), and a Finnigan LTQ orbitrap XL mass spectrometer (Thermo Scientific) equipped with an XYZ nanoelectrospray ionization source (AMR Inc.), was used for the LC/MS/MS analysis. Aliquots of trypsinized samples were automatically injected onto a peptide CapTrap cartridge (2.0 × 0.5 mm inner diameter, Michrom BioResources) attached to an injector valve for desalting and concentrating the peptides. After washing the trap with 98% H_2_O, 2% AcCN, and 0.2% TFA, the peptides were loaded into a separation capillary reverse phase column (Monocap C18 150 × 0.2 mm inner diameter, GL-Science) by switching the valve. The eluents used were as follows: A, 98% H2O, 2% AcCN, 0.1% HCOOH; and B, 10% H_2_O, 90% AcCN, 0.1% HCOOH. The column was developed at a flow rate of 1.0 μL/min, with a concentration gradient of AcCN: from 5% B to 35% B for 100 min, followed by 35% B to 95% B for one minute, maintained in 95% B for nine minutes, from 95% B to 5% B for one minute, and finally re-equilibrating with 5% B for nine minutes. Effluents were introduced into the mass spectrometer via the nanoelectrospray ion interface that held the separation column outlet directly connected with a nanoelectrospray ionization needle (PicoTip FS360–50-30; New Objective Inc.). The ESI voltage was 2.0 kV, and the transfer capillary of the LTQ inlet was heated to 200 °C. No sheath or auxiliary gas was used. The mass spectrometer was operated in a data-dependent acquisition mode, in which the MS acquisition with a mass range of m/z 420–1600 was automatically switched to MS/MS acquisition under the automated control of Xcalibur software. The top 4 precursor ions were selected by an MS scan, with Orbitrap at a resolution of *r* = 60000, and for the subsequent MS/MS scans by an ion trap in the normal/centroid mode, using the automated gain control (AGC) mode with AGC values of 5.00 × 10^5^ and 1.00 × 10^4^ for full MS and MS/MS, respectively. We also employed a dynamic exclusion capability that enabled the sequential acquisition of the MS/MS of abundant ions in the order of their intensities with an exclusion duration of two minutes and exclusion mass widths of −5 and +5 ppm. The trapping time was 100 milliseconds with the auto gain control on.

#### Reverse transcription-quantitative real-time polymerase chain reaction (RT-qPCR)

The isolated RNA was reverse transcribed into cDNA using the PrimeScript RT Master Mix (Takara). cDNA was amplified using the primer sets listed in **Table S5** with SYBR Premix Ex Taq II (Takara) in accordance with the manufacturer’s instructions. RT-qPCR analysis was performed using a Thermal Cycler Dice Real Time System (Takara). The *GAPDH* and *PGK1* mRNAs were used for transcript normalization. The relative value of the transcript was calculated using the comparative CT method (ΔΔCT). We analyzed three biological replicates with two technological replicates for each experiment.

#### Western blotting

Total protein extracts were prepared in RIPA buffer and separated on a 10% sodium dodecyl sulfate polyacrylamide gel. The proteins were transferred onto a polyvinylidene fluoride membrane (Millipore). The membrane was blocked with 3% BSA/Tris-buffered saline that contained 0.1% Triton X-100 (TBST) and was incubated with a 1:500 dilution of anti-PUM1 rabbit polyclonal antibody (Sigma-Aldrich, #HPA027424), 1:250 dilution of anti-PUM2 rabbit monoclonal antibody (Abcam, #ab92390), 1:1000 dilution of anti-GAPDH mouse monoclonal antibody (MERK, #MAB374), 1:4000 dilution of anti-FLAG M2 mouse monoclonal antibody (Sigma-Aldrich, #F1804), 1:1000 dilution of anti-CNOT1 rabbit polyclonal antibody (Protein Technology, #14276-1-AP), 1:100 dilution of anti-PCNA rabbit polyclonal antibody (Santa Cruz, #sc7907), 1:100 dilution of anti-UBE2A rabbit polyclonal antibody (Abcam, #ab31917), 1:2000 dilution of anti-Pol η rabbit polyclonal antibody (Abcam, #ab236450), or 1:1000 dilution of anti-Ub-PCNA (Lys164) rabbit monoclonal antibody (CST Technology, #13439) for one-hour at room temperature. All antibodies were diluted using IMMUNO SHOT reagent1 (COSMO BIO, Tokyo, Japan). After three washes of the membrane with TBST, the membrane was incubated with a 1:10000 or 1:5000 dilution of horseradish peroxidase (HRP)-linked secondary antibodies in IMMUNO SHOT reagent2 (COSMO BIO) for one-hour at room temperature. The proteins were visualized using Immobilon western chemiluminescent HRP substrate (MERK), according to the manufacturer’s instructions.

#### Actinomycin D chase assay

For RNA stability measurements, HeLa TO cells were treated with 1 µg/mL actinomycin D (dissolved in DMSO [Sigma-Aldrich]). After 30 minutes of incubation, the cells were harvested at the indicated time points, and RNA isolation, cDNA synthesis, and RT-qPCR were performed as previously described^79^.

#### Immunofluorescence of Pol η in cisplatin-treated cells

HCT116 cells were subcultured to 10 cm dish with 1.0×10^5^ cells/mL prior to transfection. After two days, cells were transfected with empty vector, FLAG-tagged PUM1, or FLAG-tagged ΔPUM-HD. Next day, cells were transferred to 12 well dish with 1.0×10^5^ cells/mL. Two days later, 0.2 mg/mL cisplatin (Wako) was added to the cells and incubated for 6 hr. The cells were washed with PBS and were fixed in 4% paraformaldehyde solution for 15 minutes incubation. After two times washes with PBS, cells were permeabilized with ice-cold methanol/acetone (1:1, v/v) for 10 min incubation in - 30 ℃. Then, cells were washed with PBS three times. Fixed cells were then blocked in 1%BSA/0.01% tween-20 in PBS (Blocking buffer) for 1 hour at room temperature. Cells were subsequently incubated for 1 hour at room temperature with primary antibodies (1:1000 dilution of anti-FLAG M2 antibody (Sigma-Aldrich, #F1804), 1:500 dilution of anti-Pol η antibody (Abcam, #ab236450)) in Blocking buffer. Cells were washed three times with PBS supplemented with 0.2% tween-20 and were incubated fluorophore-conjugated secondary antibodies (1:1000 dilutions of mouse-Alexa 488 and rabbit-Alexa647 (Thermo Fisher Scientific)) in Blocking buffer for one hour at room temperature. Then, cells were washed three times with PBS supplemented with 0.2% tween-20. Nuclei were counterstained with DAPI (1:1000 dilution, Wako). Fluorescent images were obtained using a fluorescence microscope (LAS AF with DMI 6000B-AFC) with HCX PL FLUOTAR L 40×/0.60na CORR PH2 objective lens (Leica). Acquired images were pseudo colored and merged using the ImageJ software. All images shown are representative of replicate experiments (*n* = 40-80).

#### BrdU incorporation assay of Pol η in cisplatin-treated cells

BrdU incorporation assay was performed using cell proliferation ELISA kit (#ab126556, Abcam) according to manufacturer’s instructions. HCT116 cells were plated in 96-well plates with 0.5×10^5^ cells/mL prior to transfection. After two days, cells were transfected with empty vector, FLAG-tagged PUM1, or FLAG-tagged ΔPUM-HD. Next day, cells were incubated for 6 hours with medium containing 0.2 mg/mL cisplatin and BrdU according to manufacturer’s recommendation. Plates were read using the Infinite F200 (TECAN, Männedorf, Switzerland) at a dual wavelength of 450/550 nm.

#### Assays for cell proliferation with cisplatin-treated cells

The proliferation of the HeLa TO cell lines was measured using the Cell Counting kit-8 (Dojindo, Kumamoto, Japan). HeLa TO cells were subcultured in 96-wells with 1.5×10^5^ cells/mL density. The next day, the cells were transfected with empty vector, FLAG-PUM1, or FLAG-ΔPUM-HD, as described in the transfection section. After 24 hours of incubation, 0.5 mg/mL of cisplatin was added to the cells for seven hours. Then, 10 μL of Cell Counting Kit-8 in 100 μL of culturing medium were added, and the cells were incubated for 30-minutes. The absorbance of 595 nm in each well was measured using Infinite F200 (TECAN, Switzerland). The mean absorption of five independent wells was calculated with the s.e.m. for each biological condition. The cell viability was defined by the absorbance of cells with cisplatin normalized by that without cisplatin.

### Quantification and Statistical Analysis

#### Preprocessing, mapping, and quantification of sequencing data

Low quality reads were discarded using the FASTX-toolkit and reads that contained less than 10 bp were eliminated using Bowtie (Langmead et al. 2009). The filtered reads were mapped to the reference human genome (hg19) and transcriptome from RefSeq downloaded from illumine iGenomes using TopHat (version 2.0.12) (Kim et al. 2013). The mapped reads were quantified to measure the RNA expression level for each gene using Cufflinks (version 2.1.1) (Trapnell et al. 2010). The abundance of RefSeq transcripts was calculated from the BRIC-seq and RIP-seq sequence reads, based on the reads per kilobase of exon model per million mapped reads (RPKM), as a means of normalizing for the gene length and depth of sequencing. Transcripts with RPKM≧ 1 were considered expressed for subsequent analysis.

#### Identification of transcripts bound to PUM

We set the threshold of the fold discovery rate, calculated using Cufflinks (version 2.1.1), to less than 0.05 for the transcripts of which the RPKM is greater than 1.0 in the corresponding RIP-PUM sample.

#### Calculation of RNA half-life and estimation of significant fold-change with PUM KD

The RNA-half life was calculated as previously described. Briefly, the relative expression data of each transcript at each time point were normalized using the 97.5% percentile. To predict an RNA decay curve, the normalized expression data were fitted to the first order exponential RNA decay model. The RNA half-life and its variance are estimated from the fitting curve and time points. In this study, the RNA half-lives were determined for 12,496 mRNAs in control cells.

To determine the significant change of the RNA half-lives, the distributions of the RNA half-lives between two conditions were compared using Welch’s *t*-test. Here, the criteria for significant stabilization were (1) log2-fold change of half-life > 1 (for de-stabilization, < 0.5) in PUM KD cells and (2) *p*-value < 0.05 (Welch’s *t*-test).

#### Database search and label-free semiquantitative proteomic analysis

MS/MS data were processed with Mascot software (version 2.5.1, Matrix Science) against the SwissProt_2016_07 database at a fragment tolerance of 0.6 Da and a parent tolerance of 5.0 ppm, considering the fixed peptide modification by carbamidomethyl (C), variable peptide modifications by methionine oxidation, and phosphorylation, including phosphoserine, phosphothreonine, and phosphotyrosine. The analysis of the data was conducted with Scaffold software (version 4.5.3, Proteome Software). The label-free semiquantitative values were determined using Expressionist Refiner MS software (GeneData). After pre-processing (noise subtraction, RT alignment, and peak detection) and importing the identification results from Mascot software, the relative ion intensities were calculated and normalized via the Lowess normalization method. The thresholds of the peptides were set to 95% minimum and the thresholds of the proteins were set to 99% minimum with 3 peptides minimum.

To determine the PUM-associated proteins, we set the threshold to a five-fold enrichment in PUM immunoprecipitants compared to the control.

#### Statistical overrepresentation test

Protein Analysis THrough Evolutionary Relationships (PANTHER) analysis was performed with 2,671 mRNAs at http://pantherdb.org/. The reference mRNA list was defined as 10,907 HeLa expressing mRNAs of which the expression is greater than 1 RPKM in RNA-seq.

#### Motif searches

The 3’UTRs of the sequences were retrieved from GenBank (RefSeq, hg19). The PUM motif was searched using MEME (http://meme-suite.org/). To assign statistical significance to the reported PUM motif, we applied the *E*-value, which is automatically calculated in the MEME program.

#### *In silico* screening of RNA-seq

To generate the complete datasets for screening, GEO was queried in terms of ‘human’, ‘inhibitor’, or ‘stress’. After the collection of the SRA files, mapping and quantification of the RNA-seq data were performed as described in the “Preprocessing, mapping, and quantification of sequencing data” section. For each RNA-seq data, the log2-fold-changes of the PUM1 and PUM1 targets were calculated. To estimate the correlation between the PUM1 and PUM1 targets, the “Variation Index of targets” was calculated as follows, *Variation Index* = “the number of increased PUM1 targets (log2-fold-change > 1)” - “the number of decreased PUM1 targets (log2-fold-change < -1)”. A scatter plot was used to investigate the potential relationship between the log2-fold-change of PUM1 and the Variation index. In the scatter plot, RNA-seq that suggests a stimulus inhibits PUM1-mediated mRNA decay will be plotted in the second quadrant, whereas RNA-seq that suggests a stimulus promotes PUM1-mediated mRNA decay will be plotted in the fourth quadrant. Clustering of log2-fold change of PUM1 target with DNA-damaging agents are made by scipy module of python. All codes are freely available at the github repository (https://github.com/Imamachi-n/AutoNGS).

#### Statistical test in q-PCR

The fold expression was calculated as 2^-ΔΔCT^. Statistical significance was defined as a p-adjusted < 0.05. For bar plots, error bars represent 1 standard deviation and asterisks (*) indicate statistically significant changes.

Data and Software Availability

#### Software

Shell scripts, python codes, and R scripts for the RNA half-life calculation are freely available at the github repository (https://github.411com/AkimitsuLab/scripts_for_BRIC-seq_data_analaysis). There are two folders named “half-life_calculation” and “half-life_calculation_for_DEG_analysis” to calculate the RNA half-life and identify the significant change of RNA half-lives between two conditions, respectively. Codes for *in silico* screening are available at https://github.com/Imamachi-n/AutoNGS.

For drawing scatter plots and a heatmap, Jupyter notebooks are available at https://github.com/ToshimichiYamada/PUM_study.

#### Data

The accession numbers for the mass spectrometry data reported in this paper are ProteomeXchange: PXD009849 (http://www.proteomexchange.org/) or JPOST: JPST000424 (https://repository.jpostdb.org/).

## Supporting Figure

**Figure S1 (related to Figure 1).**
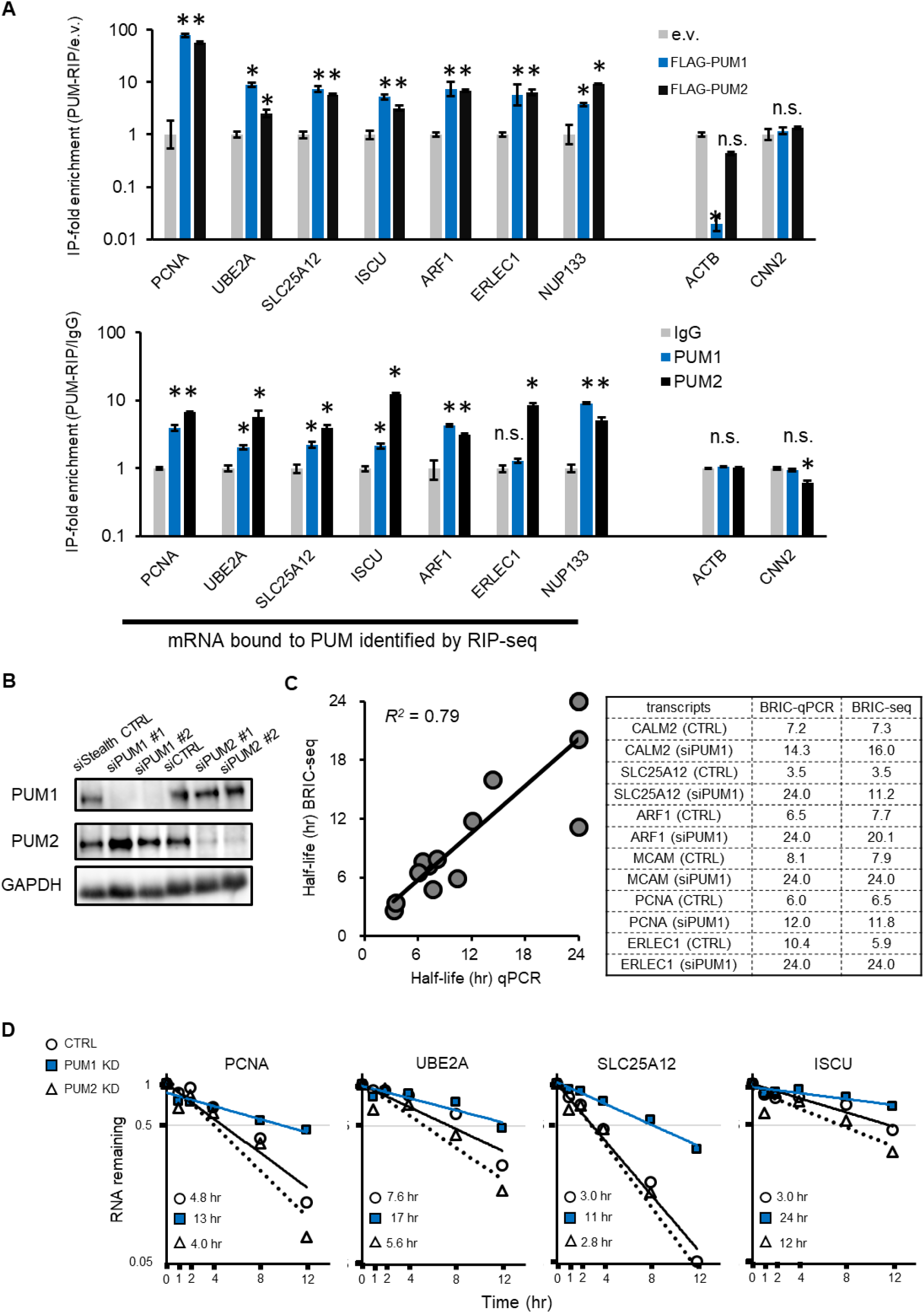
Validation of RIP-seq and BRIC-seq data by RT-qPCR. (**A**) RT-qPCR of mRNA isolated from immunoprecipitation of FLAG-tagged PUMs (upper panel) and immunoprecipitation of endogenous PUMs (lower panel) were performed to calculate the enrichment of mRNA by RIP experiments for 7 mRNAs that bound PUM and 2 non-binding mRNAs. Data are presented as the mean ± SD. n.s: not significant (*p* ≧ 0.05), *: *p* value < 0.05 (Student’s *t*-test, n=3). (**B**) Representative Western blots of PUM1, PUM2, and GAPDH from HeLa cells treated with indicated siRNA (see table S4). (**C**) A scatter plot showing the correlation of the half-lives of the indicated mRNAs determined by BRIC-seq against those calculated by RT-qPCR. Tested transcripts and values are listed in the table. (**D**) Decay curves of the indicated mRNAs determined by BRIC-seq. The half-life for each mRNA was calculated from fitted exponential functions for cells treated with a control siRNA (CTRL, circles and black line), PUM1 (blue squares and blue line), or PUM2 (triangle and black dashed line).

**Figure S2 (related to Figure 1).**
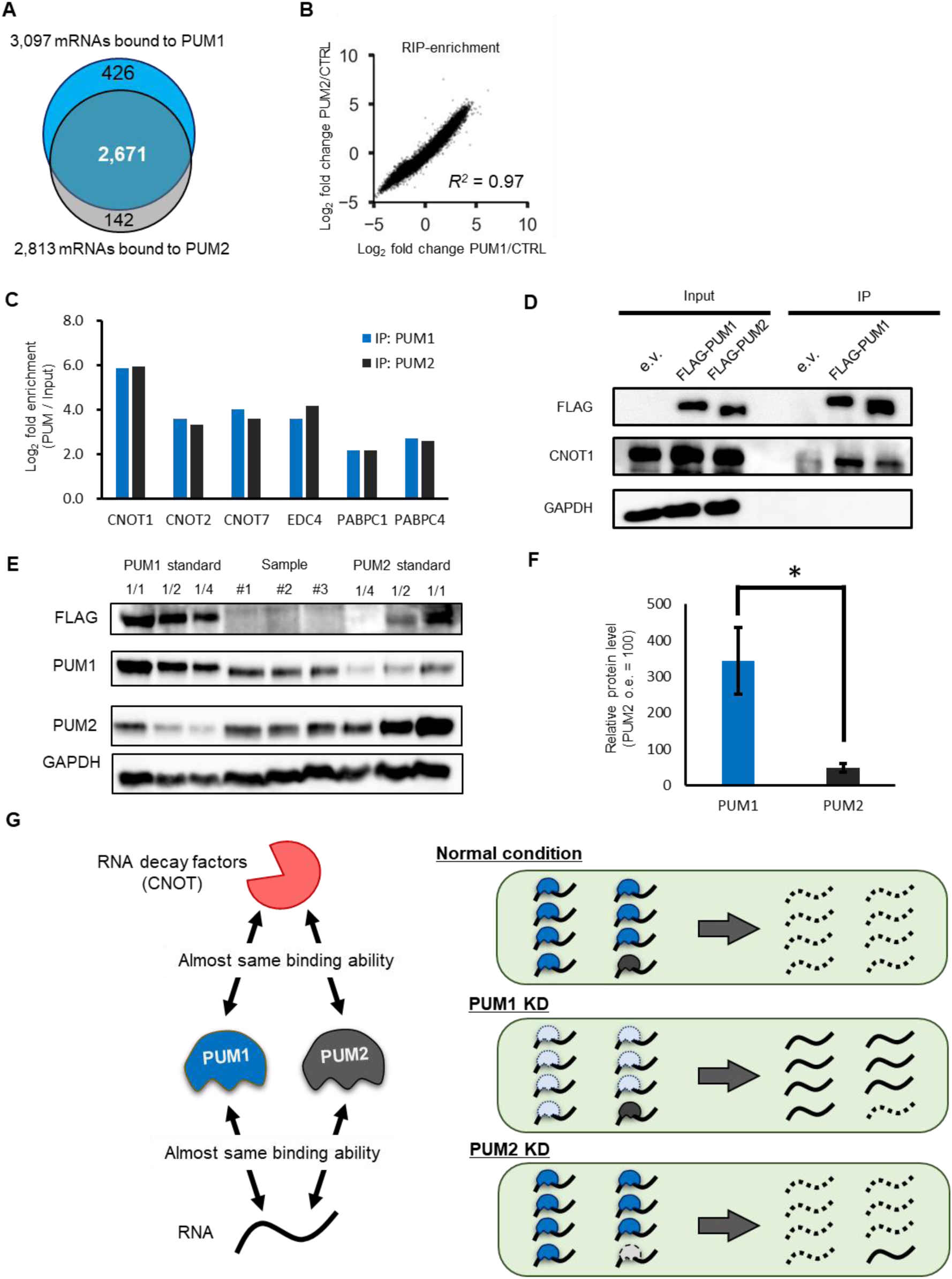
Higher abundance of PUM1 underlies the dominant effect of PUM1 on mRNA decay in HeLa TO cells. (**A**) Venn diagram showing overlap between PUM1-binding mRNAs and PUM2-binding mRNAs. (**B**) A scatter plot showing the correlation of fold enrichment levels of RIP-seq with PUM1 against RIP-seq with PUM2. (**C**) Enrichment in RNA degradation factors from mass spectrometry analysis of proteins coimmunoprecipitated with FLAG-tagged PUM1 or FLAG-tagged PUM2. The ratios of spectral counts of the indicated proteins in the PIM1 immunoprecipitates over those in immunoprecipitates of cells transfected with empty vector and the ratios for spectral counts in the PUM2 immunoprecipitates over those in immunoprecipitates from cells transfected with empty vector are shown. (See **Table S3** for list of all interacting proteins detected by mass spectrometry). (**D**) Coimmunoprecipitation of PUMs and CNOT1. The input corresponds to 20% of the total cell lysate. GAPDH served as negative control for the immunoprecipitations. (**E**) Endogenous and exogenous FLAG-PUM protein abundance analyzed by Western blot using the indicated antibodies in biological triplicates (indicated as “Sample”). GAPDH was used as a loading control for each sample. FLAG signals of exogenous FLAG-PUMs were used as a standard. Among the standard samples, 1/1, 1/2, and 1/4 indicate dilution factors. (**F**) Relative protein abundance of endogenous PUM1 proteins and PUM2 proteins. The amount of PUM2 in the cells overexpressing FLAG-PUM2 was set to 100 (PUM2 o.e. = 100). (**G**) Model of dominant PUM1-mediated mRNA decay in HeLa cells.

**Figure S3 (related to Figure 2).**
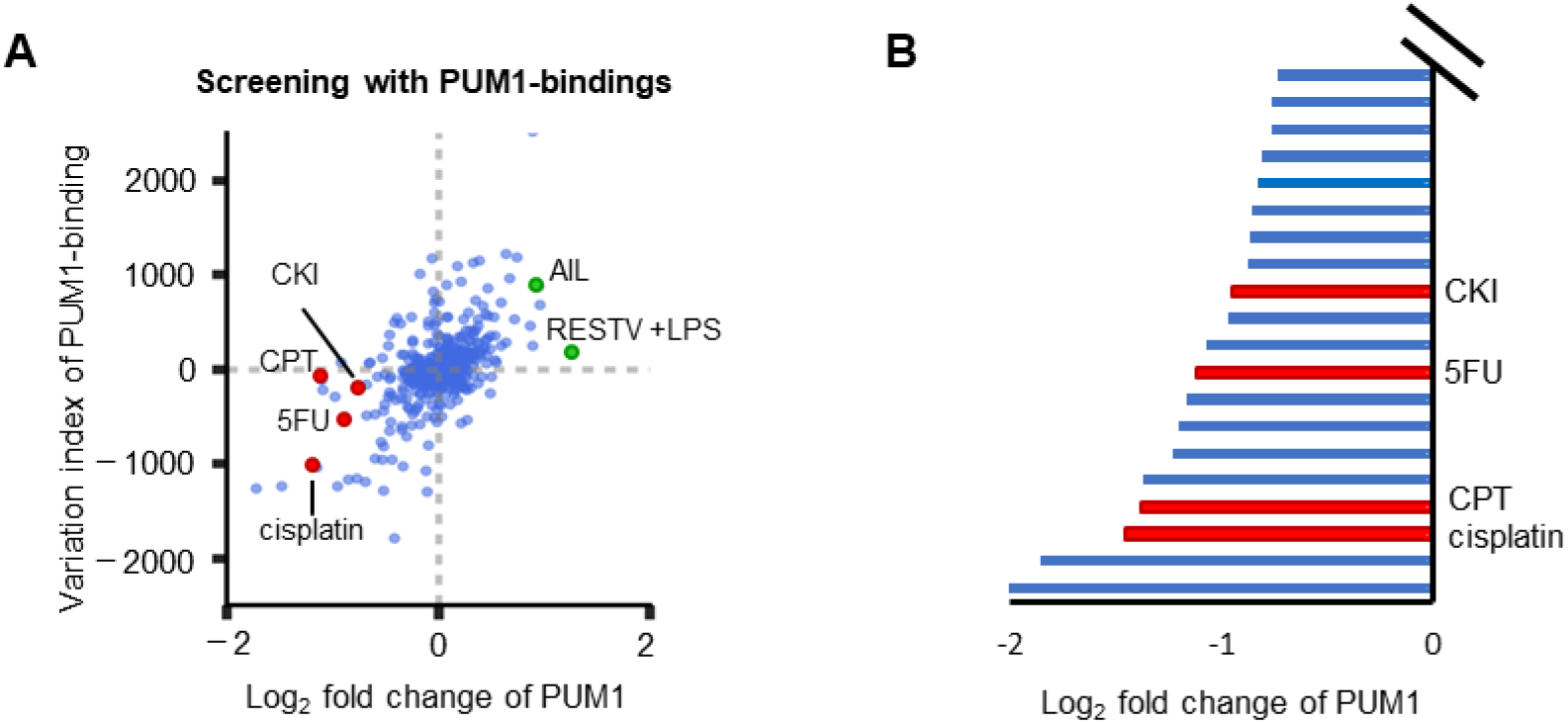
Analysis of the abundance of PUM1-binding mRNAs and PUM1 itself using complete set of PUM1-binding mRNAs. (**A**) Log_2_-fold change of PUM1 versus variation index with 491 different stimuli. Candidate inhibitory stimuli identified in Figure 2C are shown as red dots; candidate activation stimuli identified in Figure 2C are shown as green dots. Variation index was calculated from the 3,097 mRNAs that bound to PUM1 and is plotted against log_2_-fold change of PUM1. (**B**) Log_2_-fold change of PUM1 in response to various stimuli from the data in GEO. Only the response to a subset of the stimuli that includes the DNA-damaging agents highlighted as panel A are shown.

**Figure S4 (related to Figure 4).**
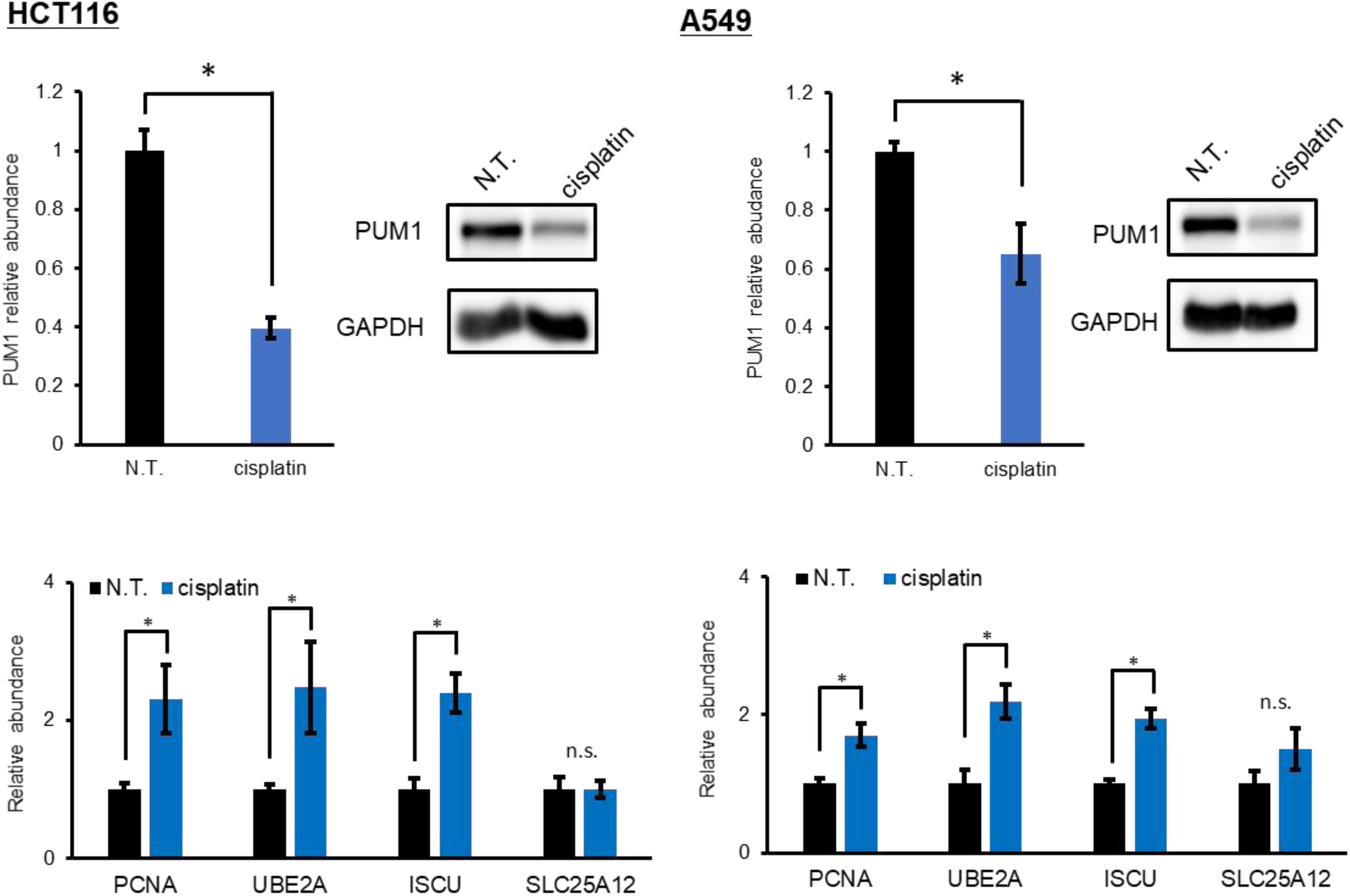
Inhibition of PUM1-mediated mRNA decay by cisplatin in HCT116 and A549 cells. (**Top**) In each cell line, Western blot analysis confirmed the reduction in PUM1 abundance. (**Bottom**) The abundance of the indicated mRNAs was determined by real-time RT-qPCR in each cell line under non-treated (N.T.) or cisplatin-treated (0.2 mg/mL for 6 hr) conditions. The abundance of each mRNA was normalized to geometric mean of *GAPDH* and *PGK1* and are shown as mean ± SD. n.s.: not significant (*p* ≧ 0.05). *: *p* value < 0.05 (Student’s *t*-test). Data are presented as mean ± SD and are from three independent experiments.

**Figure S5 (related to Figure 5).**
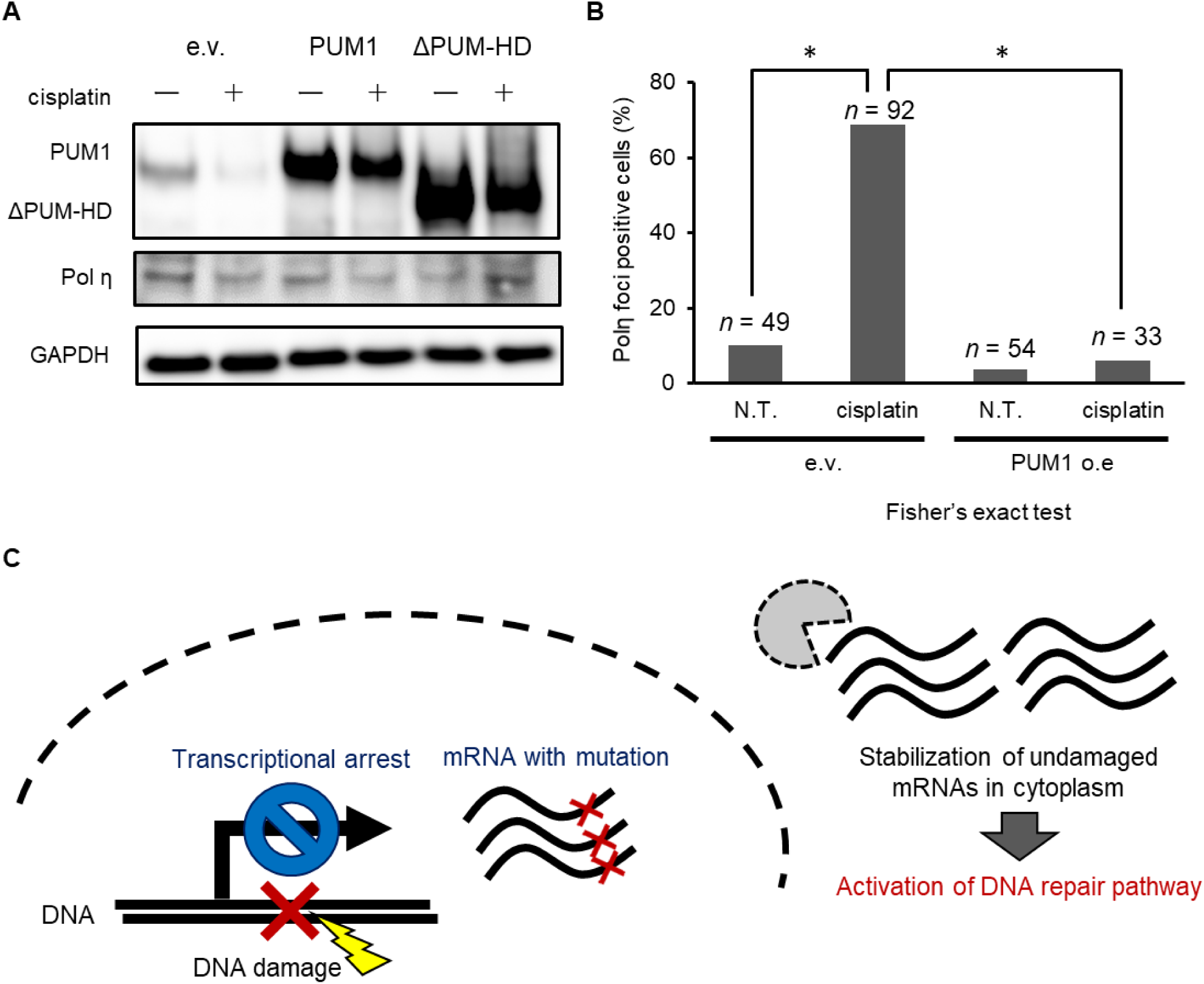
Abundance and distribution of Pol η after cisplatin treatment and a model for mRNA stabilization during DNA repair. (**A**) Western blot analysis of the Pol η from control (e.v.) or PUM1- or ΔPUM-HD-overexpressing cells with or without cisplatin (0.5 mg/mL for 6 hr) treatment. Data are representative of two experiments. (**B**) Quantification of foci-positive cells in Figure 5B. The number of cells is indicated at the top of each bar. *: *p* value < 0.05 (Fisher’s exact test). PUM1 o.e., PUM1-overexpressing cells. (**C**) Model to explain the advantage of stabilizing mRNAs in to response DNA damage.

**Table S1. mRNAs bound to and regulated by PUM1/2s** (related to **Figure 2**). This table contains the results of RIP-seq and BRIC-seq. Columns B and C show the gene symbol (symbol) and Refseq ID (refid), respectively. Half-lives (hour) of each transcript in CTRL cells (hl_CTRL) and PUM KD cells (hl_PUMKD) are indicated in Columns D and E, respectively. “nan” indicates the samples for which the half-lives were not determined because of the low accuracy of fitting. Log2-fold-change of half-life between hl_CTRL and hl_PUMKD and its *p*-value are shown as “change_hl” and “p_value” in Columns F and G, respectively. For the RIP-seq, RPKM values of input and RIP-PUM (rip_PUM1 or rip_PUM2) are shown in columns H and I, respectively. Log2-fold-change of RPKM values between input and rip_PUM and its *q*-value are illustrated as “change_rip” and “q_rip” in Columns J and K, respectively.

**Table S2. GO term enrichment analysis of PUM-bound mRNAs.** This table contains the results of the GO enrichment analysis for the biological processes. In this GO analysis, references are defined as mRNAs expressed in HeLa TO cells. Enrichment of mRNAs bound to PUM were analyzed against the reference. GO terms with FDR (column H) less than 0.05 are shown.

**Table S3. List of proteins interacting with PUM1/2** (related to **Figure S2**). This table summarizes the proteins that interact with Flag-tagged PUM1 or PUM2. Column A shows the gene symbol. Molecular weight of each protein is indicated in column B. Spectral counts between immunoprecipitated (IPed) fragment with e.v. (IP-e.v.), PUM1 (IP-PUM1), or PUM2 (IP-PUM2) are listed in columns C to E. The ratio of spectral counts (IP-PUM1/IP-PUM2, IP-PUM1/IP-e.v., and IP-PUM2/IP-e.v.) are shown in columns F-H. If the value of IP-e.v. is zero, we transformed the value as 0.5 to calculate the ratio.

**Table S4. List of RNA-seq used for *in silico* screening** (related to **Figures 3 and 6**). IDs (Bioproject ID, GEO ID, and SRA ID) of each RNA-seq are obtained from GEO. From the information for cell lines and stimulating conditions, we manually curated the RNA-seq and defined the control sample (C#) and treated sample (T#) to calculate the log2-fold change of RNA expression with stimulus.

**Table S5. List and sequence of siRNAs and primers for RT-qPCR.**

## References

1. Nadal, E. de, Ammerer, G. & Posas, F. Controlling gene expression in response to stress. Nat. Rev. Genet. 12, 833–845 (2011).

2. Rabani, M. et al. Metabolic labeling of RNA uncovers principles of RNA production and degradation dynamics in mammalian cells. Nat. Biotechnol. 29, 436–442 (2011).

3. Rabani, M. et al. High-Resolution Sequencing and Modeling Identifies Distinct Dynamic RNA Regulatory Strategies. Cell 159, 1698–1710 (2014).

4. Schoenberg, D. R. & Maquat, L. E. Regulation of cytoplasmic mRNA decay. Nat. Rev. Genet. 13, 246–259 (2012).

5. Chen, C.-Y. A. & Shyu, A.-B. Mechanisms of deadenylation-dependent decay. Wiley Interdiscip. Rev. RNA 2, 167–183 (2011).

6. Collart, M. A. The Ccr4-Not complex is a key regulator of eukaryotic gene expression. Wiley Interdiscip. Rev. RNA 7, 438–454 (2016).

7. Müller-McNicoll, M. & Neugebauer, K. M. How cells get the message: dynamic assembly and function of mRNA–protein complexes. Nat. Rev. Genet. 14, 275–287 (2013).

8. Dominguez, D. et al. Sequence, Structure, and Context Preferences of Human RNA Binding Proteins. Mol. Cell 70, 854–867.e9 (2018).

9. Wheeler, E. C., Van Nostrand, E. L. & Yeo, G. W. Advances and challenges in the detection of transcriptome-wide protein-RNA interactions. Wiley Interdiscip. Rev. RNA 9, e1436 (2018).

10. Nussbacher, J. K. & Yeo, G. W. Systematic Discovery of RNA Binding Proteins that Regulate MicroRNA Levels. Mol. Cell 69, 1005–1016.e7 (2018).

11. Imamachi, N., Salam, K. A., Suzuki, Y. & Akimitsu, N. A GC-rich sequence feature in the 3′ UTR directs UPF1-dependent mRNA decay in mammalian cells. Genome Res. 27, 407–418 (2016).

12. Wang, M., Ogé, L., Perez-Garcia, M.-D., Hamama, L. & Sakr, S. The PUF Protein Family: Overview on PUF RNA Targets, Biological Functions, and Post Transcriptional Regulation. Int. J. Mol. Sci. 19, 410 (2018).

13. Bohn, J. A. et al. Identification of diverse target RNAs that are functionally regulated by human Pumilio proteins. Nucleic Acids Res. 46, 362–386 (2018).

14. Wreden, C., Verrotti, A. C., Schisa, J. A., Lieberfarb, M. E. & Strickland, S. Nanos and pumilio establish embryonic polarity in Drosophila by promoting posterior deadenylation of hunchback mRNA. Development 124, 3015–23 (1997).

15. Joly, W., Chartier, A., Rojas-Rios, P., Busseau, I. & Simonelig, M. The CCR4 Deadenylase Acts with Nanos and Pumilio in the Fine-Tuning of Mei-P26 Expression to Promote Germline Stem Cell Self-Renewal. Stem Cell Reports 1, 411–424 (2013).

16. Gennarino, V. A. et al. Pumilio1 Haploinsufficiency Leads to SCA1-like Neurodegeneration by Increasing Wild-Type Ataxin1 Levels. Cell 160, 1087–1098 (2015).

17. Driscoll, H. E., Muraro, N. I., He, M. & Baines, R. A. Pumilio-2 Regulates Translation of Nav1.6 to Mediate Homeostasis of Membrane Excitability. J. Neurosci. 33, 9644–9654 (2013).

18. Xu, E. Y., Chang, R., Salmon, N. A. & Reijo Pera, R. A. A gene trap mutation of a murine homolog of theDrosophila stem cell factorPumilio results in smaller testes but does not affect litter size or fertility. Mol. Reprod. Dev. 74, 912–921 (2007).

19. Mak, W., Fang, C., Holden, T., Dratver, M. B. & Lin, H. An Important Role of Pumilio 1 in Regulating the Development of the Mammalian Female Germline. Biol. Reprod. 94, 134 (2016).

20. Spassov, D. S. & Jurecic, R. Cloning and comparative sequence analysis of PUM1 and PUM2 genes, human members of the Pumilio family of RNA-binding proteins. Gene 299, 195–204 (2002).

21. Goldstrohm, A. C., Hall, T. M. T. & McKenney, K. M. Post-transcriptional Regulatory Functions of Mammalian Pumilio Proteins. Trends Genet. 34, 972–990 (2018).

22. Galgano, A. et al. Comparative Analysis of mRNA Targets for Human PUF-Family Proteins Suggests Extensive Interaction with the miRNA Regulatory System. PLoS One 3, e3164 (2008).

23. Hafner, M. et al. Transcriptome-wide Identification of RNA-Binding Protein and MicroRNA Target Sites by PAR-CLIP. Cell 141, 129–141 (2010).

24. Morris, A. R., Mukherjee, N. & Keene, J. D. Ribonomic Analysis of Human Pum1 Reveals cis-trans Conservation across Species despite Evolution of Diverse mRNA Target Sets. Mol. Cell. Biol. 28, 4093–4103 (2008).

25. Etten, J. Van et al. Human Pumilio Proteins Recruit Multiple Deadenylases to Efficiently Repress Messenger RNAs. J. Biol. Chem. 287, 36370–36383 (2012).

26. Weidmann, C. A., Raynard, N. A., Blewett, N. H., Van Etten, J. & Goldstrohm, A. C. The RNA binding domain of Pumilio antagonizes poly-adenosine binding protein and accelerates deadenylation. RNA 20, 1298–1319 (2014).

27. Goldstrohm, A. C., Hook, B. A., Seay, D. J. & Wickens, M. PUF proteins bind Pop2p to regulate messenger RNAs. Nat. Struct. Mol. Biol. 13, 533–539 (2006).

28. Webster, M. W., Stowell, J. A. & Passmore, L. A. RNA-binding proteins distinguish between similar sequence motifs to promote targeted deadenylation by Ccr4-Not. Elife 8, (2019).

29. Campbell, Z. T. et al. Cooperativity in RNA-protein interactions: global analysis of RNA binding specificity. Cell Rep. 1, 570–81 (2012).

30. Jarmoskaite, I. et al. A Quantitative and Predictive Model for RNA Binding by Human Pumilio Proteins. Mol. Cell (2019). doi:10.1016/j.molcel.2019.04.012

31. Hu, Z., Killion, P. J. & Iyer, V. R. Genetic reconstruction of a functional transcriptional regulatory network. Nat. Genet. 39, 683–687 (2007).

32. Bar-Joseph, Z. et al. Computational discovery of gene modules and regulatory networks. Nat. Biotechnol. 21, 1337–1342 (2003).

33. Lee, T. I. et al. Transcriptional Regulatory Networks in Saccharomyces cerevisiae. Science (80-.). 298, 799–804 (2002).

34. Muhar, M. et al. SLAM-seq defines direct gene-regulatory functions of the BRD4-MYC axis. Science (80-.). 360, 800–805 (2018).

35. Yamada, T. & Akimitsu, N. Contributions of regulated transcription and mRNA decay to the dynamics of gene expression. Wiley Interdiscip. Rev. RNA 10, e1508 (2019).

36. Zhao, J. et al. Genome-wide Identification of Polycomb-Associated RNAs by RIP-seq. Mol. Cell 40, 939–953 (2010).

37. Tani, H. et al. Genome-wide determination of RNA stability reveals hundreds of short-lived noncoding transcripts in mammals. Genome Res. 22, 947–956 (2012).

38. Yamada, T. et al. 5′-Bromouridine IP Chase (BRIC)-Seq to Determine RNA Half-Lives. In *Methods in molecular biology (Clifton*, N.J.) 1720, 1–13 (2018).

39. Jakymiw, A. et al. Disruption of GW bodies impairs mammalian RNA interference. Nat. Cell Biol. 7, 1267–1274 (2005).

40. Nishi, K., Nishi, A., Nagasawa, T. & Ui-Tei, K. Human TNRC6A is an Argonaute-navigator protein for microRNA-mediated gene silencing in the nucleus. RNA 19, 17–35 (2013).

41. Elkayam, E. et al. Multivalent Recruitment of Human Argonaute by GW182. Mol. Cell 67, 646–658.e3 (2017).

42. Lee, S. et al. Noncoding RNA NORAD Regulates Genomic Stability by Sequestering PUMILIO Proteins. Cell 164, 69–80 (2016).

43. Wang, W. et al. Anti-tumor activities of active ingredients in Compound Kushen Injection. Acta Pharmacol. Sin. 36, aps201524 (2015).

44. Qu, Z. et al. Identification of candidate anti-cancer molecular mechanisms of compound kushen injection using functional genomics. Oncotarget 7, 66003–66019 (2016).

45. Aguilera, A. & Gómez-González, B. Genome instability: a mechanistic view of its causes and consequences. Nat. Rev. Genet. 9, 204–217 (2008).

46. Basu, A. & Krishnamurthy, S. Cellular Responses to Cisplatin-Induced DNA Damage. J. Nucleic Acids 2010, 1–16 (2010).

47. Friedberg, E. C., Lehmann, A. R. & Fuchs, R. P. P. Trading Places: How Do DNA Polymerases Switch during Translesion DNA Synthesis? Mol. Cell 18, 499–505 (2005).

48. Kawamoto, T. et al. Dual Roles for DNA Polymerase η in Homologous DNA Recombination and Translesion DNA Synthesis. Mol. Cell 20, 793–799 (2005).

49. Masutani, C., Kusumoto, R., Iwai, S. & Hanaoka, F. Mechanisms of accurate translesion synthesis by human DNA polymerase eta. EMBO J. 19, 3100–9 (2000).

50. Somasagara, R. R. et al. RAD6 promotes DNA repair and stem cell signaling in ovarian cancer and is a promising therapeutic target to prevent and treat acquired chemoresistance. Oncogene 36, 6680–6690 (2017).

51. Sanders, M. A., Haynes, B., Nangia-Makker, P., Polin, L. A. & Shekhar, M. P. Pharmacological targeting of RAD6 enzyme-mediated translesion synthesis overcomes resistance to platinum-based drugs. J. Biol. Chem. 292, 10347–10363 (2017).

52. Hoege, C., Pfander, B., Moldovan, G.-L., Pyrowolakis, G. & Jentsch, S. RAD6-dependent DNA repair is linked to modification of PCNA by ubiquitin and SUMO. Nature 419, 135–141 (2002).

53. Bienko, M. et al. Regulation of Translesion Synthesis DNA Polymerase η by Monoubiquitination. Mol. Cell 37, 396–407 (2010).

54. Sekimoto, T., Oda, T., Kurashima, K., Hanaoka, F. & Yamashita, T. Both High-Fidelity Replicative and Low-Fidelity Y-Family Polymerases Are Involved in DNA Rereplication. Mol. Cell. Biol. 35, 699–715 (2015).

55. Kannouche, P. et al. Localization of DNA polymerases eta and iota to the replication machinery is tightly co-ordinated in human cells. EMBO J. 21, 6246–56 (2002).

56. Kannouche, P. et al. Domain structure, localization, and function of DNA polymerase eta, defective in xeroderma pigmentosum variant cells. Genes Dev. 15, 158–72 (2001).

57. Roos, W. P. & Kaina, B. DNA damage-induced cell death by apoptosis. Trends Mol. Med. 12, 440–450 (2006).

58. Roos, W. P. & Kaina, B. DNA damage-induced cell death: From specific DNA lesions to the DNA damage response and apoptosis. Cancer Lett. 332, 237–248 (2013).

59. Lieberthal, W., Triaca, V. & Levine, J. Mechanisms of death induced by cisplatin in proximal tubular epithelial cells: apoptosis vs. necrosis. Am. J. Physiol. Physiol. 270, F700–F708 (1996).

60. Kannouche, P. L., Wing, J. & Lehmann, A. R. Interaction of human DNA polymerase eta with monoubiquitinated PCNA: a possible mechanism for the polymerase switch in response to DNA damage. Mol. Cell 14, 491–500 (2004).

61. Blattner, C. et al. UV-Induced stabilization of c-fos and other short-lived mRNAs. Mol. Cell. Biol. 20, 3616–25 (2000).

62. Wang, W. et al. HuR regulates p21 mRNA stabilization by UV light. Mol. Cell. Biol. 20, 760– 9 (2000).

63. Prives, C. & Gottifredi, V. The p21 and PCNA partnership: A new twist for an old plot. Cell Cycle 7, 3840–3846 (2008).

64. Soria, G., Speroni, J., Podhajcer, O. L., Prives, C. & Gottifredi, V. p21 differentially regulates DNA replication and DNA-repair-associated processes after UV irradiation. J. Cell Sci. 121, 3271–3282 (2008).

65. Tichon, A. et al. A conserved abundant cytoplasmic long noncoding RNA modulates repression by Pumilio proteins in human cells. Nat. Commun. 7, 12209 (2016).

66. Lee, S. et al. Noncoding RNA NORAD Regulates Genomic Stability by Sequestering PUMILIO Proteins. Cell 164, 69–80 (2016).

67. Munschauer, M. et al. The NORAD lncRNA assembles a topoisomerase complex critical for genome stability. Nature 561, 132–136 (2018).

68. Wickens, M., Bernstein, D. S., Kimble, J. & Parker, R. A PUF family portrait: 3′UTR regulation as a way of life. Trends Genet. 18, 150–157 (2002).

69. Gerber, A. P., Herschlag, D. & Brown, P. O. Extensive Association of Functionally and Cytotopically Related mRNAs with Puf Family RNA-Binding Proteins in Yeast. PLoS Biol. 2, e79 (2004).

70. Quenault, T., Lithgow, T. & Traven, A. PUF proteins: repression, activation and mRNA localization. Trends Cell Biol. 21, 104–112 (2011).

71. Kaye, J. A., Rose, N. C., Goldsworthy, B., Goga, A. & L’Etoile, N. D. A 3’UTR pumilio-binding element directs translational activation in olfactory sensory neurons. Neuron 61, 57– 70 (2009).

72. Kedde, M. et al. A Pumilio-induced RNA structure switch in p27-3′ UTR controls miR-221 and miR-222 accessibility. Nat. Cell Biol. 12, 1014–1020 (2010).

73. Chen, D. et al. Pumilio 1 Suppresses Multiple Activators of p53 to Safeguard Spermatogenesis. Curr. Biol. 22, 420–425 (2012).

74. Mukhopadhyay, R. et al. DAPK-ZIPK-L13a Axis Constitutes a Negative-Feedback Module Regulating Inflammatory Gene Expression. Mol. Cell 32, 371–382 (2008).

75. Arif, A. et al. Two-Site Phosphorylation of EPRS Coordinates Multimodal Regulation of Noncanonical Translational Control Activity. Mol. Cell 35, 164–180 (2009).

76. Sundararaman, B. et al. Resources for the Comprehensive Discovery of Functional RNA Elements. Mol. Cell 61, 903–913 (2016).

77. Lambert, N. et al. RNA Bind-n-Seq: Quantitative Assessment of the Sequence and Structural Binding Specificity of RNA Binding Proteins. Mol. Cell 54, 887–900 (2014).

78. Nostrand, E. L. Van et al. Robust transcriptome-wide discovery of RNA-binding protein binding sites with enhanced CLIP (eCLIP). Nat. Methods 13, nmeth.3810 (2016).

79. Yang, E. et al. Decay Rates of Human mRNAs: Correlation With Functional Characteristics and Sequence Attributes TL - 13. Genome Res. 13 VN-r, 1863 (2003).

